# DNA supercoiling restricts the transcriptional bursting of neighboring eukaryotic genes

**DOI:** 10.1101/2022.03.04.482969

**Authors:** Heta P. Patel, Stefano Coppola, Wim Pomp, Ineke Brouwer, Tineke L. Lenstra

## Abstract

DNA supercoiling has emerged as a major contributor to gene regulation in bacteria. The impact of DNA supercoiling on transcription dynamics in eukaryotes is less clear. Here, using single-molecule dual-color RNA imaging in budding yeast, we show that transcriptional bursting of the divergent and tandem *GAL* genes is coupled. Upon topoisomerase degradation, supercoils that buildup from transcription inhibit subsequent transcription at neighboring genes, thereby reducing their simultaneous bursting. *GAL* gene transcription is inhibited more by negative than by positive supercoiling accumulation. Unlike bacteria, wildtype yeast has sufficient topoisomerase levels to minimize inhibition from supercoils at adjacent genes. Overall, we discover fundamental differences in supercoiling-mediated gene regulation between bacteria and yeast and show that rapid supercoiling release in eukaryotes ensures proper gene expression of neighboring genes.

**One sentence summary:** Transcription causes twisting of the DNA double helix, which can inhibit transcription of adjacent genes.

## INTRODUCTION

Transcription, the process of copying DNA to RNA, is an essential process in all living cells. During transcription, movement of RNA polymerase along the DNA generates negative supercoils behind RNA polymerase and positive supercoils in front, as described by the twin-supercoiled-domain model (*1*–*3*). Transcription-generated supercoils can, in turn, enhance or impede the transcriptional process: negative supercoils facilitate transcription initiation by enabling promoter melting and enhancing the binding of regulatory factors, whereas positive supercoils aid the elongation of RNA polymerases by destabilizing DNA-bound proteins (*4*–*9*). However, excessive negative or positive supercoils can also repress transcription (*10*–*14*). The impact of supercoiling-mediated inhibition is illustrated in *E. coli*, which have a limited concentration of the topoisomerase gyrase. At highly transcribed genes, the dynamic accumulation and release of positive supercoils stochastically switch genes off and on, thereby causing transcriptional bursting (*10*). Whether eukaryotic topoisomerase levels are limiting and how positive and negative DNA supercoiling control transcription dynamics in eukaryotes is still unknown.

Transcription-generated supercoils can propagate along the DNA and may activate or deactivate adjacent genes. For multiple bacterial species, negative supercoils generated behind polymerase enhance transcription of upstream divergent genes, whereas positive supercoils in front of polymerase inhibit transcription of downstream tandem and convergent genes (*15*–*19*). Similar mechanisms were proposed in eukaryotes, but direct *in vivo* evidence is lacking (*20*–*26*). In contrast to bacterial DNA, eukaryotic DNA is wrapped in nucleosomes, which may buffer excess positive supercoils to limit their dissipation (*27, 28*). However, chromatin does not absorb negative supercoils (*29*). Accordingly, negative supercoils propagate up to 1.5kb around the transcription start site of transcribed genes (*23*). Whether *in vivo* negative supercoils enhance transcription of divergent genes and whether positive supercoils are efficiently buffered by nucleosomes to prevent the inhibition of tandem and convergent genes remains unclear.

Supercoiling-mapping studies observed that gene bodies are positively supercoiled, and promoters are negatively supercoiled (*23, 30*–*32*). Gene promoters are maintained in a negatively supercoiled state by restricting the activity of topoisomerase TOP1 to gene bodies (*33*). In addition, mammalian genomes contain large negatively-supercoiled domains of actively transcribed genes (*32*). However, since mapping of supercoils at the single-cell level is technically challenging, little is known about the differences in supercoiling states between cells at a single time point, or how supercoiling dynamics affect the transcription dynamics of single eukaryotic cells over time.

In this study, we used the closely-positioned and highly-expressed divergent (*GAL1-GAL10*) and tandem (*GAL10-GAL7*) gene pairs of the *GAL* gene cluster in *S. cerevisiae* to investigate how DNA supercoiling affects transcriptional bursting of neighboring eukaryotic genes (**Figure 1A**). Using single-molecule dual-color imaging, we found that transcriptional bursting of the *GAL* gene pairs was temporally correlated inside single cells and that yeast topoisomerases are essential for maintaining the correlation. At the divergent *GAL1*-*GAL10* genes, excess negative supercoils impede, rather than enhance, transcription. At the tandem *GAL10-GAL7* genes, in addition to positive supercoils inhibiting transcription of the downstream gene, we discover that negative supercoils considerably inhibit transcription of the upstream gene. Moreover, wildtype budding yeast has sufficient concentrations of topoisomerases to minimize the inhibition of DNA supercoiling on transcription, implying that DNA supercoils play different regulatory roles on gene transcription in prokaryotes versus eukaryotes.

**Fig. 1:**
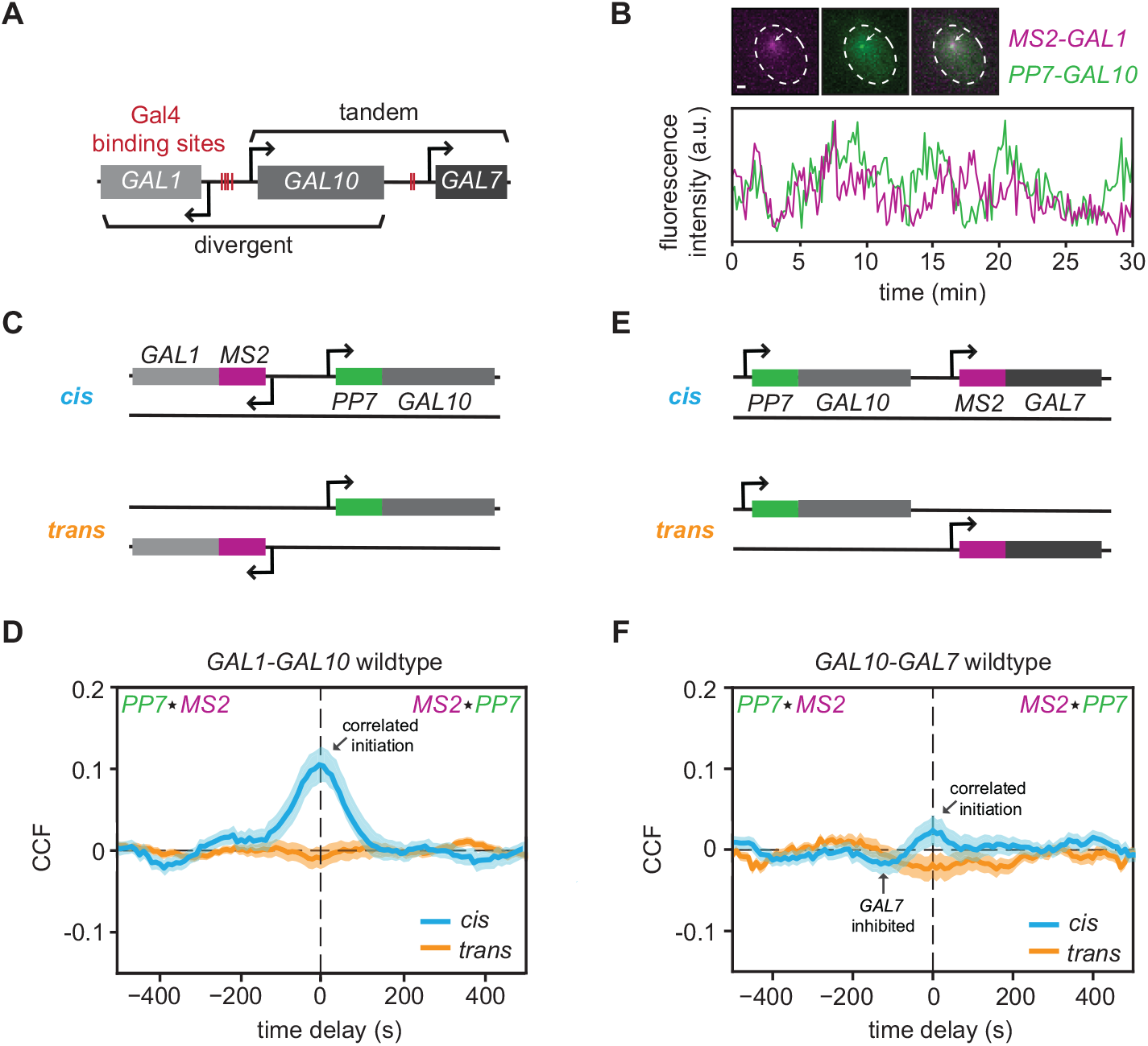
Transcriptional bursting of the divergent and tandem *GAL* genes is temporally coupled. **(A)** Schematic of *GAL* gene cluster in yeast. Red lines indicate the binding sites of the transcription factor, Gal4. The gene lengths of *GAL1, GAL10* and *GAL7* are approximately 1.5 kb, 2.1 kb, 1.1 kb, respectively. The intergenic distance of the divergent *GAL1-GAL10* genes is 669 bp and of the tandem *GAL10-GAL7* genes is 726 bp. All three genes are highly transcriptionally active in galactose-containing media and therefore produce high levels of DNA supercoils at the locus (*30*). In galactose, the antisense transcripts at this locus are not transcribed (*73*). **(B)** Example images of *MS2*-*GAL1* (magenta), *PP7*-*GAL10* (green) and merged (gray) transcription sites (TSs), indicated by arrows (top). Scale bar: 1 µm. Example traces of the quantified fluorescence intensities (arbitrary units) of the *MS2*-*GAL1* and *PP7*-*GAL10* TSs (bottom). **(C)** Nascent transcription of the divergent *GAL1-GAL10* gene pair is visualized either on the same allele (*cis*) or different alleles (*trans*). **(D)** *MS2-PP7* cross-correlation of the divergent *GAL1-GAL10* genes in the *cis* configuration (blue) and the *trans* control (orange). *cis*: *n* = 179 cells; *trans*: *n* = 197 cells. Shaded area indicates SEM. **(E)** Same as (**C**) for the tandem *GAL10-GAL7* genes. **(F)** Same as (**D**) for the tandem *GAL10-GAL7* genes. *cis*: *n* = 148 cells; *trans*: *n* = 125 cells.

## RESULTS

### Transcriptional bursting of the divergent and tandem *GAL* genes is temporally coupled

To visualize nascent transcription at the divergent *GAL* gene pair with single-molecule resolution (**Figure 1B**), we inserted 12 repeats of MS2V6 and 14 repeats of PP7 at the 5’ untranslated regions of *GAL1* and *GAL10*, respectively (**Figure 1C**) (*34, 35*). Upon transcription, these repeats form loops that are specifically bound by the fluorescently-tagged MS2 and PP7 coat proteins, allowing for nascent RNA visualization at the endogenous loci in living cells. The divergent genes were labeled on the same chromosome (*cis* configuration) or on two different chromosomes (*trans* configuration) to distinguish between local environment effects and extrinsic noise effects, such as correlations generated by cell-to-cell variations.

To determine if the transcriptional bursting of the divergent *GAL1-GAL10* genes was temporally coupled, we imaged live cells that were induced with galactose, quantified the intensities of the *MS2-GAL1* and *PP7-GAL10* transcription sites (TSs), and computed the *MS2-PP7* cross-correlation function (CCF) (**Figures 1B, D, S1A-B**). The CCF measures the similarity of *MS2-GAL1* and *PP7-GAL10* time traces at varying time delays. The *MS2-PP7* CCF of the *cis*-labeled genes displayed a defined peak at time delay zero, indicating that *GAL1* and *GAL10* initiate together (**Figure 1D**). The CCF decayed to zero at time delays of approximately -100s and +100s, which was in accordance with the decay of *GAL1* and *GAL10* from the auto-correlation functions (ACFs) (**Figure S1E**). The CCF of the *trans*-labeled divergent genes yielded a flat line, as expected for independently-expressed, uncorrelated genes on different chromosomes. The magnitude of the correlation at zero time delay was quantified using the normalized transcriptional overlap (**Methods**) (*36*), which represents the percentage of co-occurring *GAL1-GAL10* transcription events when *GAL10* is active. Because of the high activity of these genes, we observed substantial transcriptional overlap (75±2%) for the *trans* control. The *cis* configuration exhibited a higher overlap (85±2%), demonstrating that the transcription of the divergent genes is more correlated in the *cis* configuration than in the *trans* (**Figure S1F**).

These results were corroborated using single-molecule RNA fluorescence *in situ* hybridization (smFISH) with probes that hybridized to the MS2 and PP7 loops (*37*). As a measure for correlated transcription, we computed the Pearson correlation coefficient of the number of nascent RNA transcripts at the *MS2-GAL1* and *PP7-GAL10* TSs across thousands of cells (**Figure S1G**). Consistent with the live-cell results, *GAL1*-*GAL10* transcription shows a higher Pearson correlation in the *cis* (R=0.20±0.02) than in the *trans* (R=0.12±0.02) configuration. Therefore, when positioned on the same chromosome, the divergent *GAL1* and *GAL10* genes initiate simultaneously, more than by random chance.

Next, the same approach was used to measure the transcriptional correlation of the tandem *GAL10*-*GAL7* genes (**Figures 1E, S1C-E**). The *cis*-labeled tandem genes are weakly correlated at time delay zero (**Figure 1F**) with a transcriptional overlap that is 2.0±0.3% higher compared to the *trans* control (**Figure S1H**). The modest positive correlation difference could not be confirmed by smFISH (**Figure S1I**), presumably because it is obscured by extrinsic noise. We also observed a weak valley in the CCF at time delay -100s, which implies that 100s subsequent to a burst of *GAL10, GAL7* transcription events are observed less than by random chance. Because of the well-established inhibitory role of positive supercoils, we expect that positive supercoils generated from *GAL10* elongation may inhibit *GAL7* initiation (*10, 19, 31*). Such periods of lower transcription initiation rates immediately after a transcriptional burst have been referred to as refractory periods (*38*–*40*). In this manuscript, we employ the same nomenclature, regardless of whether the refractory period follows transcription of the gene itself or of the neighboring gene. Overall, live-cell imaging and smFISH indicate that transcription of the divergent and tandem *GAL* gene pairs in wildtype yeast is temporally coupled.

### Conditional degradation of topoisomerases results in refractory periods

In yeast, transcription-generated supercoiling levels are managed by topoisomerases, Top1 and Top2, which act redundantly to relieve both positive and negative supercoils (*41*). To investigate how DNA supercoiling affects the temporal correlation of the divergent and tandem *GAL* gene pairs, we perturbed DNA supercoiling levels by conditionally degrading endogenous Top1 and Top2 using the auxin-inducible degron system (*42*). The presence of OsTIR1, without the addition of auxin, resulted in basal degradation of degron-tagged Top1 and Top2 (**Figure S2A-B**). Basal degradation was exploited by expressing one or two copies of OsTIR1, resulting in 25% and 40% degradation, respectively. Upon addition of auxin, Top1 and Top2 were efficiently degraded within 15 min (**Figure S2A-B**). This strategy yielded a range of yeast strains with varying levels of topoisomerases to measure the effects of supercoiling accumulation on transcription.

Live-cell imaging of *GAL1*-*GAL10* and *GAL10*-*GAL7* transcription in these strains revealed that topoisomerase degradation introduced valleys in the CCF around -100s and +100s time delays **(Figures 2, S2C-H)**. These valleys indicate that 100s after a correlated burst, simultaneous transcription of the two genes is observed less often than expected by random chance. A transcriptional burst of *GAL1*, either alone or together with *GAL10*, causes a refractory period for *GAL10* and vice versa. For a few conditions, asymmetric valleys in the CCFs illustrate that one gene is inhibited more than the other (**Figure 2B, F**). Specifically, for *GAL1-GAL10*, the inhibition was stronger for *GAL10*, while for *GAL10-GAL7*, the inhibition was stronger for *GAL7*. Similar but weaker valleys at 100s time delay were observed in the ACFs, most prominently for *GAL10*, indicating *GAL10* transcription may also weakly inhibit itself (**Figure S2I-T**). Finally, the presence of recurring peaks every 200s in the CCF indicates periodicity, likely resulting from the refractory period. Taken together, these results suggest a model where the accumulation of DNA supercoils from a transcriptional burst limits subsequent transcription of neighboring genes.

**Fig. 2:**
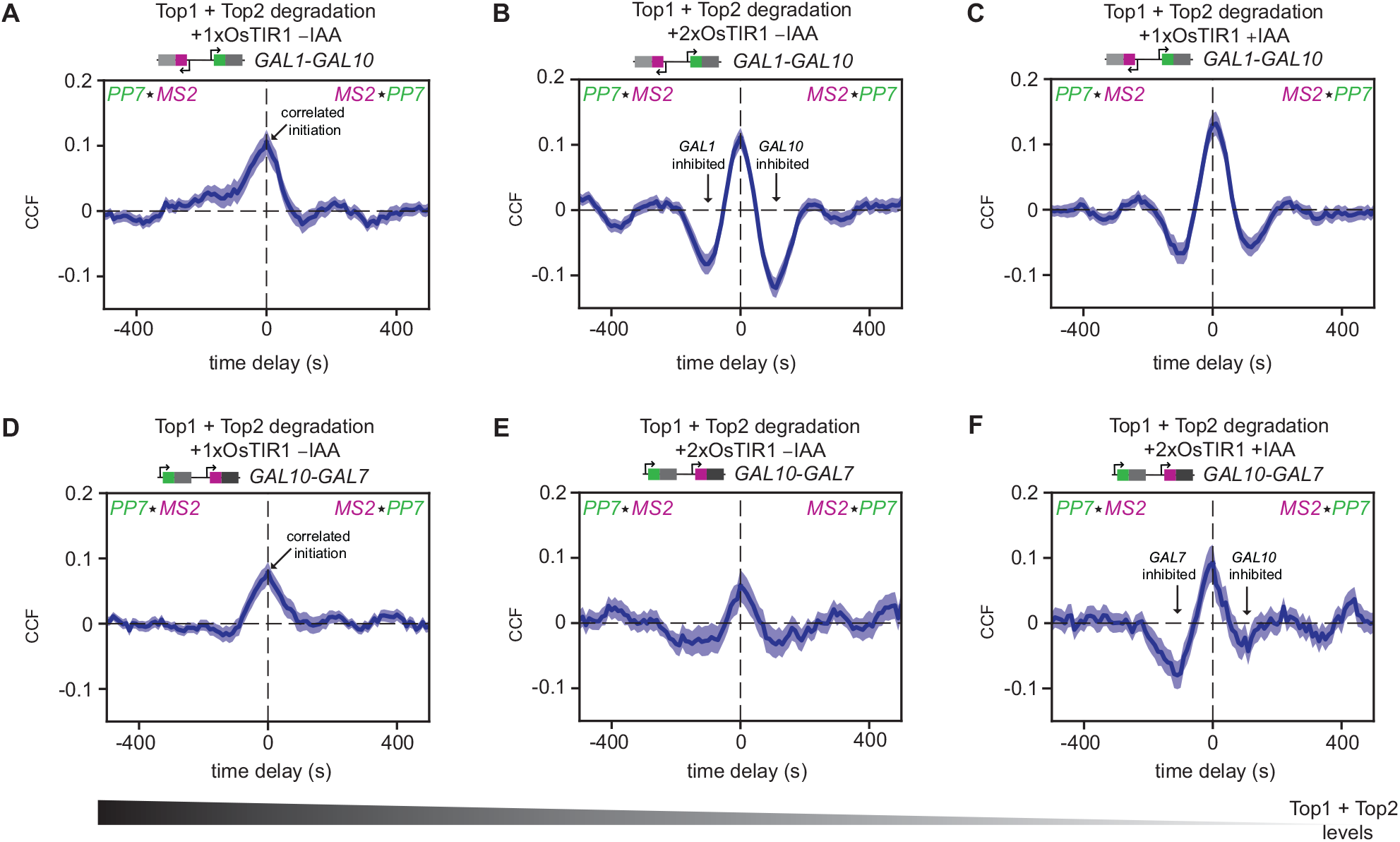
Conditional degradation of topoisomerases results in refractory periods. (**A-C**) *MS2-PP7* cross-correlation of the divergent *GAL1-GAL10* genes in cells with indicated OsTIR1 and auxin conditions. Arrows indicate examples of peaks and valleys. *n* = 117 cells, 214 cells, and 273 cells, respectively. Shaded area indicates SEM. (**D-F**) Same as (**A-C**) for the tandem *GAL10-GAL7* genes. *n* = 188 cells, 119 cells, and 143 cells, respectively.

### Conditional degradation of topoisomerases reduces the simultaneous initiation of neighboring *GAL* genes

The refractory period at neighboring genes upon supercoiling accumulation suggests that these genes initiate together less often than in wildtype. In agreement, quantification of the transcriptional overlap at zero time delay of *GAL1-GAL10* and *GAL10-GAL7* upon topoisomerase degradation revealed a reduction in the overlap for both gene pairs (**Figure 3A-B**). We note that the transcriptional overlap is normalized to the transcriptional activity of the genes. smFISH experiments confirmed that decreasing topoisomerase levels progressively decreased the *GAL1-GAL10* and *GAL10-GAL7* Pearson correlation coefficients of transcriptionally active cells (**Figure 3C-F**). The decreased correlation for *GAL1-GAL10* and *GAL10-GAL7* gene pairs was also corroborated in untagged wildtype strains using gene-specific smFISH probes (**Figure S3A-B**). The basal Top1 and Top2 degradation and its associated phenotype could partially be rescued by the addition of the antagonist, auxinole, confirming the specificity of the effects (**Figure S3C-N**). These results indicate that the divergent and tandem *GAL* genes initiate together less frequently when supercoils accumulate at the locus. For the divergent *GAL1-GAL10* genes, this reduced correlation challenges previous models, which predicted that in eukaryotes, the accumulation of negative supercoils in divergent promoters enhances the correlation of divergent genes (*22*–*25, 43*).

**Fig. 3:**
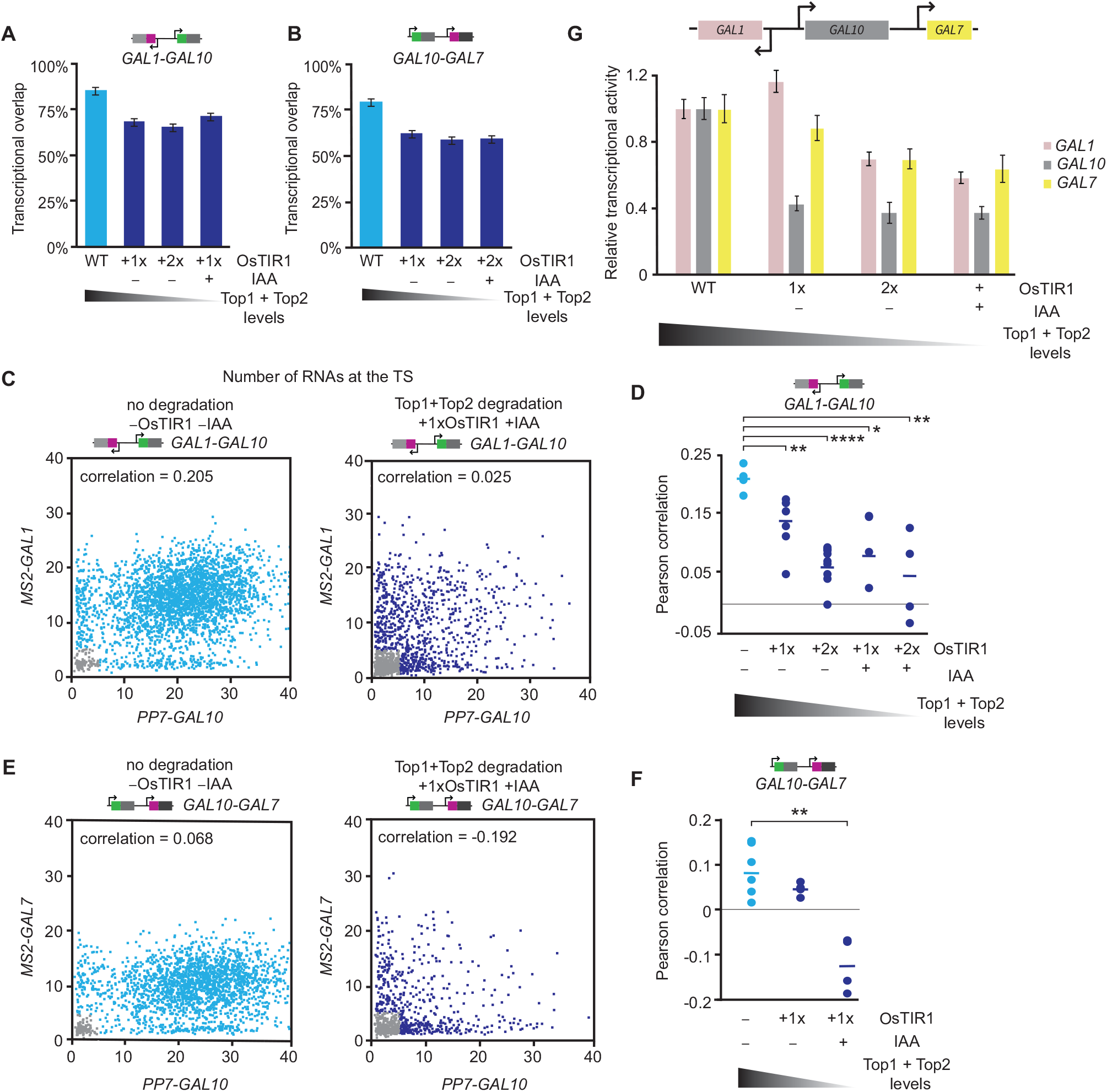
Conditional degradation of topoisomerases reduces the simultaneous initiation of neighboring genes. (**A-B**) Transcriptional overlap of WT (blue) and topoisomerase-deficient cells (navy) for the (A) divergent *GAL1-GAL10* and (B) tandem *GAL10-GAL7* gene pairs. Error bars indicate SEM. **(C)** Scatterplots of the number of nascent transcripts at the *MS2-GAL1* and *PP7-GAL10* TSs, determined by smFISH, in WT (left, *n* = 9,529 cells) and topoisomerase-depleted (right, *n* = 1,411 cells) cells. Each datapoint represents a cell. Gray datapoints represent transcriptionally inactive cells that were excluded from the Pearson correlation coefficient calculation. **(D)** Pearson correlation coefficients of *GAL1-GAL10* nascent transcription from smFISH in WT (blue) and topoisomerase-deficient strains (navy). Each circle represents a single smFISH replicate experiment (*n* = 4, 6, 7, 3, 4, from left to right). Horizontal lines represent means. All replicates consist of at least 500 cells. \**p*<0.05; \*\**p*<0.01; \*\*\*\**p*<0.0001, determined by two-tailed t-test. **(E)** Same as (**C**), for *MS2-GAL7* and *PP7-GAL10* in WT (left, *n* = 2,865 cells) and topoisomerase-deficient cells (right, *n* = 940 cells). **(F)** Same as (**D**), for the tandem *GAL10-GAL7* genes (*n* = 5, 3, 3). \*\**p*<0.01, determined by two-tailed t-test. **(G)** Relative transcriptional activity of *GAL1, GAL10* and *GAL7* in WT and topoisomerase-deficient cells, calculated by taking the inverse of the ACF amplitudes. WT is normalized to 1. Error bars indicate SEM.

To understand why supercoiling accumulation reduced the correlated transcription of the *GAL* gene pairs, we analyzed the relative transcriptional activity of the three *GAL* genes from the inverse amplitude of the ACFs. Complete topoisomerase degradation reduced the transcriptional activity of all three genes, while partial topoisomerases degradation inhibited *GAL10* transcription more than *GAL1* and *GAL7* (**Figure 3G**). Analysis of the bursting parameters using binarized *MS2* and *PP7* intensity traces revealed that topoisomerase inhibition results in shorter-duration, lower-intensity, and lower-frequency bursts for all genes, with the largest effect at *GAL10* (**Figure S2U-W**). The uneven inhibition of *GAL10* is supported by smFISH, in which the percentage of actively transcribing cells is reduced more for *GAL10* than for *GAL1* (**Figure S2X-Y**). The ACF of *GAL10* also showed a prominent refractory period, which was not as evident for *GAL1* and *GAL7* (**Figure S2I-T**). The specific inhibition of *GAL10* may explain the loss in correlation of *GAL1-GAL10* and *GAL10-GAL7* upon supercoiling accumulation. Overall, we conclude that topoisomerases are important for maintaining high transcription levels and ensuring correlated transcription between the divergent and tandem *GAL* genes.

### Transcription-generated supercoils from *GAL7* inhibit *GAL10* transcription in topoisomerase-deficient conditions

We hypothesized that the disproportionate inhibition of *GAL10* transcription in topoisomerase-deficient cells (**Figure 3G**) is caused by the accumulation of transcription-generated DNA supercoils from the highly expressed downstream gene *GAL7*. To test this hypothesis, we used two complementary approaches to limit the possible effects of *GAL7*-generated supercoils (**Figure 4A-B**).

**Fig. 4:**
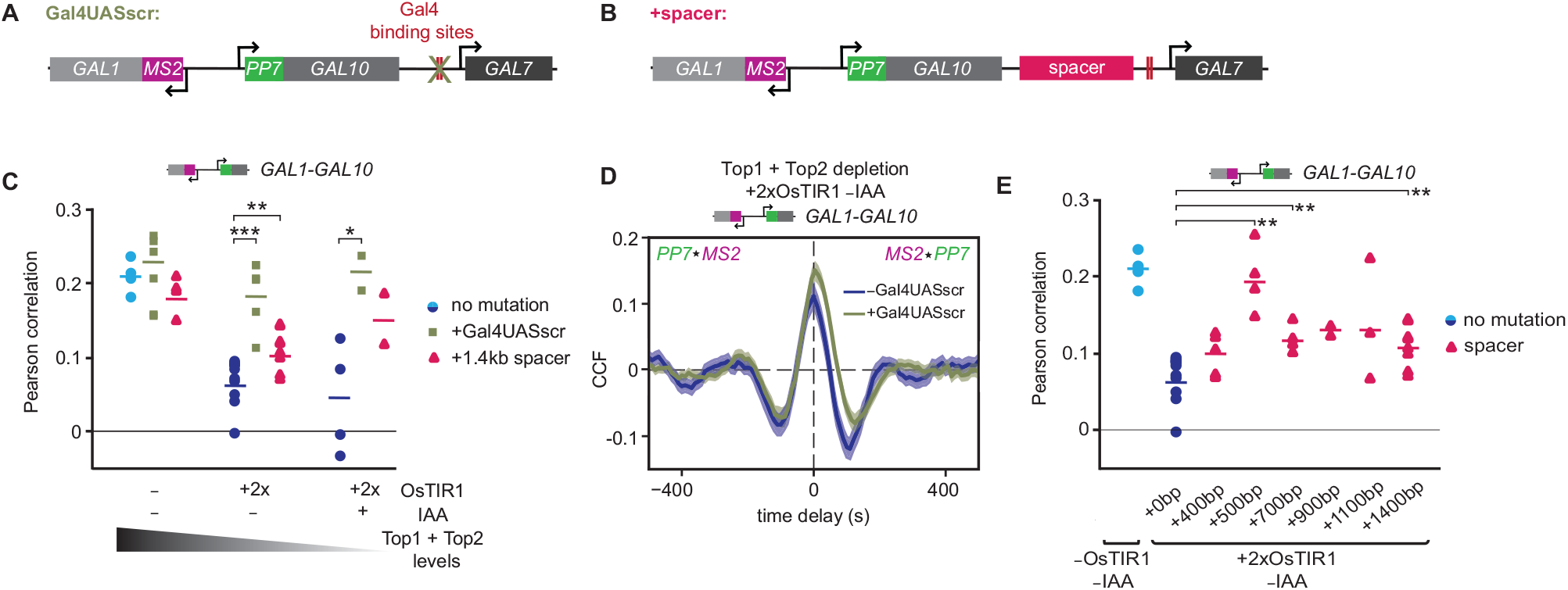
Transcription-generated supercoils from *GAL7* inhibit *GAL10* transcription in topoisomerase-deficient conditions. (**A-B**) Schematic of the *GAL* locus with: (A) scrambled Gal4UAS sites (Gal4UASscr) in the *GAL7* promoter and (B) insertion of a spacer sequence in the *GAL10-GAL7* intergenic region. **(C)** Pearson correlation coefficients of *GAL1-GAL10* nascent transcription from smFISH of WT (blue circles) and topoisomerase-deficient cells (navy circles) with Gal4UASscr (green squares) and insertion of a 1.4kb spacer (magenta triangles). Horizontal lines represent mean. Each symbol represents a single smFISH replicate experiment (*n* = 4, 5, 4, 7, 4, 6, 4, 2, 2, from left to right). All experiments consist of at least 500 cells. \**p*<0.05; \*\**p*<0.01; \*\*\**p*<0.001, determined by two-tailed t-test. **(D)** Overlay of *MS2-PP7* cross-correlation of *GAL1-GAL10* in topoisomerase-deficient cells with Gal4UASscr (green) and with wildtype Gal4UAS (navy; same as **Figure 2C**). Shaded area indicates SEM. **(E)** Same as (**C**) for *GAL1-GAL10* with increasing spacer sequence lengths (pink triangles) (*n* = 4, 7, 5, 4, 4, 2, 3, 6). \*\**p*<0.01, determined by two-tailed t-test.

First, transcription of *GAL7* was abolished by scrambling both Gal4 upstream activating sequences (Gal4 UAS) in the *GAL7* promoter (**Figures 4A, S4A**). We expected that the lack of DNA supercoils from *GAL7* transcription in this mutant would increase *GAL10* expression and therefore increase the correlation between *GAL1* and *GAL10*. As predicted, smFISH of topoisomerase-deficient cells showed an increase in the fraction of cells transcribing *GAL10* (**Figure S4F**) and a rescue in the *GAL1-GAL10* correlation to wildtype levels (**Figures 4C**). Live-cell imaging of the divergent *GAL1-GAL10* genes upon loss of *GAL7* transcription showed dampened valleys at +100s time delay and loss of periodicity in the *GAL10* ACF and *GAL1-GAL10* CCF, revealing a weaker *GAL10* refractory period (**Figures 4D, S4B-E**). *GAL7* inhibition also partially rescued the *GAL1-GAL10* transcriptional overlap at zero time delay (**Figure S4C**). These results demonstrate that the transcription-generated DNA supercoils from *GAL7* inhibit *GAL10*. Interestingly, in wildtype cells, elimination of *GAL7* transcription did not affect the fraction of cells transcribing *GAL10* (**Figure S4F**), nor the *GAL1-GAL10* correlation (**Figure 4C**), suggesting that wildtype cells possess sufficient topoisomerase levels to prevent the inhibition of *GAL7*-generated supercoils on *GAL10* transcription.

As a second method to test whether *GAL7*-generated DNA supercoils inhibit *GAL10* transcription, a 1.4 kb spacer sequence was inserted between *GAL10* and *GAL7* to dissipate transcription-generated supercoils (**Figure 4B, C**). In topoisomerase-deficient cells, addition of a spacer increased the *GAL10* active fraction and partially rescued the correlation of the divergent genes, corroborating supercoiling-mediated inhibition of *GAL10* (**Figures 4C, S4G**). Similar to the Gal4 binding site perturbation, addition of the spacer did not affect the correlation of the divergent genes in wildtype cells (**Figure 4C**). Moreover, insertion of spacers with various lengths revealed an optimal intergenic distance of 500 bp that fully rescues the *GAL1-GAL10* correlation with partial rescues at other distances (**Figure 4E**). The reason for this optimal distance is unclear. Overall, these perturbations demonstrate that transcription-generated supercoils from *GAL7* inhibit *GAL10* in topoisomerase-deficient cells but not in wildtype cells.

### Transcription inhibition at the *GAL* locus is predominantly caused by negative supercoils

Our experiments indicate *GAL10* and *GAL7* mutually inhibit each other (**Figures 1F, 2, 4**). The tandem *GAL10-GAL7* orientation suggests that an accumulation of positive supercoils from the *GAL10* gene body inhibits *GAL7* (**Figures 1F, 2D-F**), whereas an accumulation of negative supercoils from the *GAL7* promoter inhibits *GAL10* (**Figure 4**). To distinguish the contribution of positive or negative supercoils on transcription inhibition, we ectopically overexpressed bacterial topoisomerases gyrase or DNA topoisomerase I (Topo I), which selectively relieve positive or negative supercoils, respectively, and have been shown to work in yeast (*30, 44, 45*). Relieving excess positive supercoils with ectopic gyrase expression in topoisomerase-deficient cells weakly increased the correlation of the divergent *GAL1*-*GAL10* genes (**Figure 5A**), but weakly decreased the correlation of the tandem *GAL10-GAL7* genes (**Figure 5B**). The modest effects on transcription confirm that positive supercoils in eukaryotes are efficiently buffered by nucleosomes or relieved by topoisomerases (*27, 46*).

**Fig. 5:**
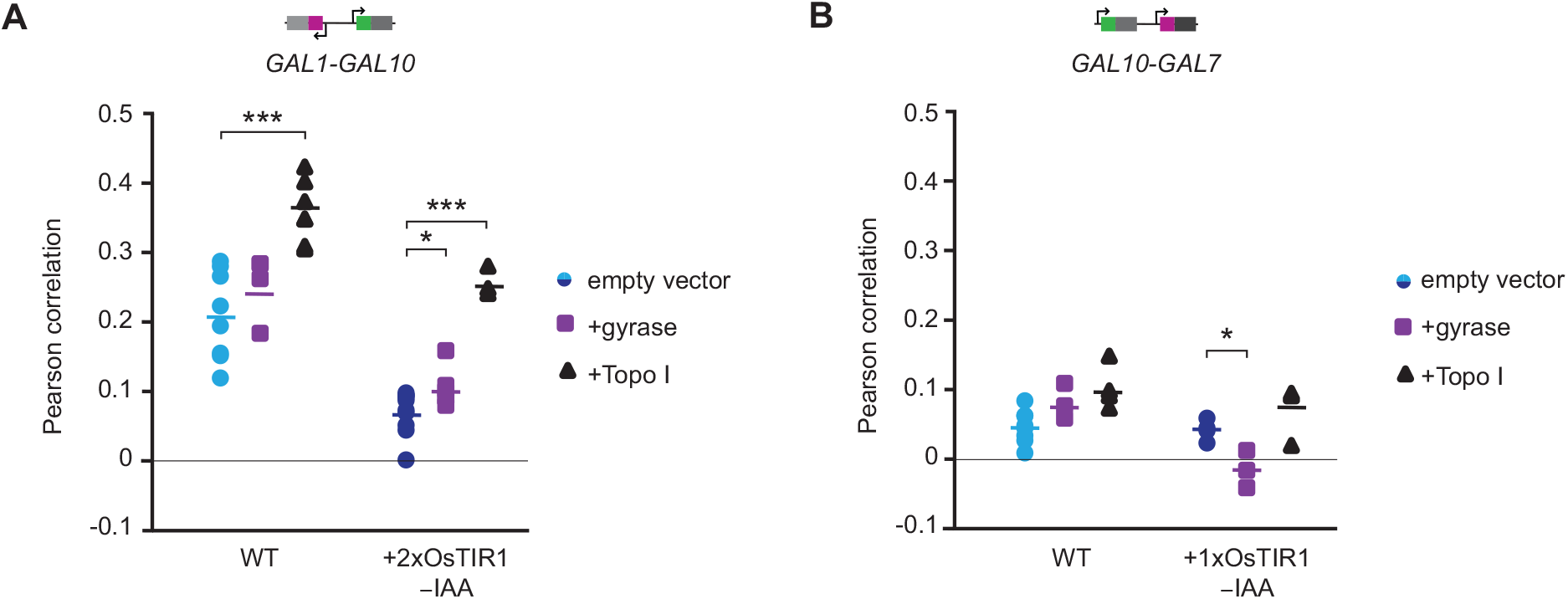
Transcription inhibition at the *GAL* locus is predominantly caused by negative supercoils. **(A)** Pearson correlation coefficients of *GAL1-GAL10* nascent transcription by smFISH with ectopic expression of gyrase (purple squares) and Topo I (black triangles) in WT and topoisomerase-deficient cells. Horizontal lines represent mean. Each symbol represents a single smFISH replicate experiment (*n* = 7, 3, 6, 6, 4, 3, from left to right). All experiments consist of at least 500 cells. \*\*\**p*<0.001, determined by two-tailed t-test. **(B)** Same as (**A**) for tandem *GAL10-GAL7* genes (*n* = 6, 4, 4, 3, 3, 3). \**p*<0.05, determined by two-tailed t-test.

Overexpression of bacterial Topo I to relax excess negative supercoils considerably increased the correlation of the divergent pair (**Figure 5A**). A small subpopulation exhibits higher expression of *GAL1* and *GAL10* than wildtype, partly explaining the increased *GAL1-GAL10* correlation (**Figure S5C**). For the tandem *GAL10-GAL7* genes, Topo I overexpression did not significantly change the correlation (**Figure 5B**). In these overexpression experiments, the timing of Topo I induction coincides with the timing of the observed effects on the correlations, indicating that these observations are unlikely to be caused by indirect effects (**Figure S5A-B**). We conclude that at the *GAL* locus, transcription is considerably more inhibited by an accumulation of negative than positive supercoils.

### Supercoiling-mediated inhibition is not caused by altered chromatin structure

Previous studies have shown that an accumulation of supercoils can change nucleosome stability (*7, 9*). To gain mechanistic insight into the supercoiling-mediated transcription inhibition, we tested if the accumulation of DNA supercoils influenced the position and stability of nucleosomes at the *GAL* locus. We performed micrococcal nuclease digestion, followed by sequencing (MNase-seq) in wildtype, basal topoisomerase degradation (1xOsTIR -IAA) and full degradation (1xOsTIR +IAA) conditions (**Figure S6A-B**) using high and low concentrations of MNase to map stable and fragile nucleosomes, respectively (*47*).

In basal topoisomerase degradation conditions, we observed that the position of stable and fragile nucleosomes in the promoters of the divergent and tandem *GAL* gene pairs is unchanged (**Figures 6A, S6C**), even though transcription inhibition was observed in these conditions. Only at full topoisomerase degradation did minor shifts in the nucleosome position appear at the locus. Similarly, genome-wide analysis showed that only full degradation of topoisomerases resulted in less well-positioned stable nucleosomes (**Figure 6B**). However, since these effects were not observed upon basal topoisomerase degradation, we conclude that changed nucleosome positioning or stability is not the main cause of the observed transcription inhibition upon supercoiling accumulation and instead, may be an independent effect of supercoiling accumulation.

**Fig. 6:**
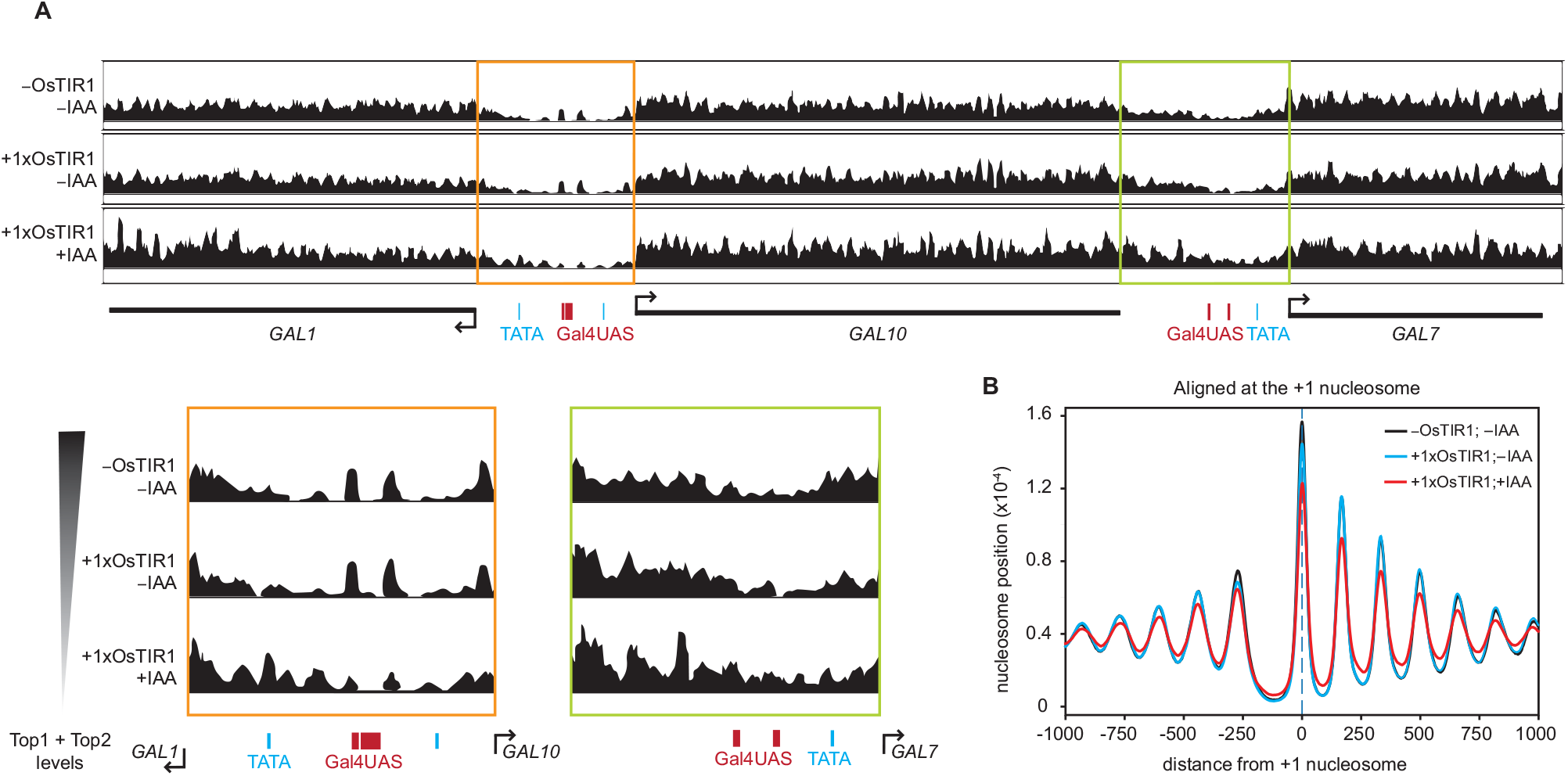
Supercoiling-mediated inhibition is not caused by altered chromatin structure. **(A)** MNase-seq profiles depicting the stable nucleosome midpoint positions at the *GAL* locus in WT (top) and topoisomerase-deficient (middle, bottom) haploid cells. Midpoints are smoothed by 31 bp. Zoom-in of the *GAL1-GAL10* and *GAL10-GAL7* promoter regions depicted in orange and green boxes, respectively. Positions of Gal4 binding sites (red) and TATA boxes (blue) are indicated. **(B)** Average nucleosome positions of all genes, aligned at the +1 nucleosome, for WT (black) and topoisomerase-deficient cells (blue, red).

### Supercoiling accumulation does not alter Gal4 and TBP binding

We next investigated whether the accumulation of supercoils affect the ability of the transcription factor and TATA-binding protein (TBP) to bind to the DNA, as has been suggested previously (*13, 14*). Using chromatin immunoprecipitation followed by qPCR (ChIP-qPCR), we measured the occupancy of the transcription factor Gal4 at the Gal4 UASs, and of TBP at the TATA boxes in the *GAL1* and *GAL10* promoters. In contrast to previous experiments with temperature sensitive mutants (*14*), we found that neither basal nor full degradation of topoisomerases reduced Gal4 and TBP occupancies (**Figure S6D-F**). Therefore, supercoiling-mediated inhibition is not caused by altered Gal4 or TBP binding but is likely caused by a step subsequent to TBP binding.

### DNA supercoils inhibit tandem genes genome-wide

To understand if the supercoiling-mediated inhibition is also observed in population-based genome-wide measurements, we used a published yeast Rpb1 ChIP-seq dataset, and analyzed the fold change in polymerase occupancy at gene pairs upon topoisomerase degradation (*48*). All adjacent and annotated gene pairs were categorized based on their relative orientations (tandem, convergent, and divergent) and for tandem gene pairs, we performed the analysis for upstream and downstream genes separately. Within each category, gene pairs were binned based on the transcription levels of the neighboring gene (**Figure 7**). Since absolute ChIP occupancies are difficult to quantitatively compare between samples without spike-in normalization, our analysis focused on relative differences between highly- and lowly-expressed gene pairs. We expected that supercoils generated from high transcription levels inhibit the transcription of neighboring genes upon topoisomerase degradation.

**Fig. 7:**
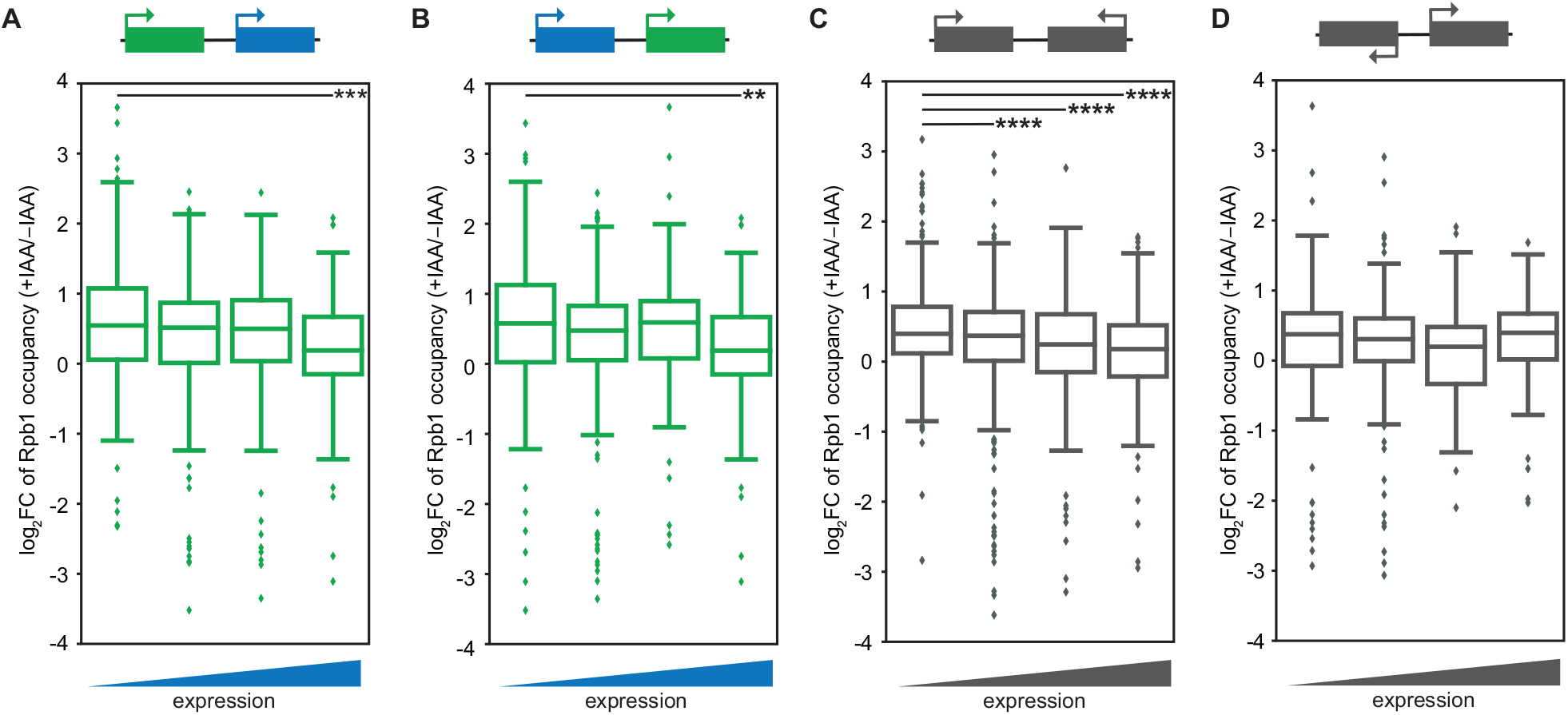
DNA supercoils inhibit tandem genes genome-wide. **(A)** Boxplot of the log_2_ fold change in Rpb1 occupancy upon topoisomerase degradation of upstream tandem genes (green) with varying expression of downstream genes (blue). Low = 87 genes; medium = 135 genes; high = 36 genes; very high = 29 genes. The box indicates quartiles, the horizontal line inside the box indicates the median, and the whiskers indicate 1.5 times the interquartile range of the box. Data points outside the whiskers are indicated with diamonds. \*\*\**p*<0.001; determined by two-tailed t-test and Bonferroni correction for multiple (three comparisons) test. **(B)** Same as (**A**) for downstream tandem genes (green) with varying expression of upstream genes (blue). Low = 171 genes; medium = 384 genes; high = 128 genes; very high = 71 genes. \*\**p*<0.01; determined by two-tailed t-test and Bonferroni correction for multiple (three comparisons) test. **(C)** Boxploht of the log_2_ fold change in Rpb1 occupancy upon topoisomerase degradation of convergent genes. Low = 372 genes; medium = 696 genes; high = 163 genes; very high = 124 genes. \*\*\*\**p*<0.0001; determined by two-tailed t-test and Bonferroni correction for multiple (three comparisons) test. **(D)** Same as (**C**) for divergent gene pairs, where the neighboring gene has varied expression. Low = 267 genes; medium = 531 genes; high = 159 genes; very high = 100 genes.

Consistent with our prediction, integrated Pol II occupancy of a gene decreased more upon topoisomerase degradation when a tandem neighboring gene was highly expressed (**Figure 7A-B)**. Similar to *GAL10-GAL7*, tandem gene pairs genome-wide were mutually repressed by an upstream gene as well as a downstream gene (**Figure 7A-B**), presumably by the accumulation of positive and negative supercoils, respectively. We also observed a transcription-dependent downregulation at convergent gene pairs, in accordance with the inhibitory effect of positive supercoils (**Figure 7C**). However, inhibition from supercoiling accumulation is not observed for divergent genes in these population-based measurements (**Figure 7D**), underscoring the importance of time-resolved experiments to detect temporally-restricted transcription patterns. Overall, we conclude that the supercoiling-mediated inhibition observed at *GAL10*-*GAL7* occurs at tandem genes genome-wide.

## Discussion

In this study, we combined single-molecule transcription imaging at neighboring *GAL* genes with targeted perturbations to expose how transcription-generated DNA supercoiling shapes transcription dynamics in budding yeast. We find that DNA supercoiling accumulation causes a temporally restricted expression pattern, where transcription initiation of a gene can occur simultaneously with its neighbor, but initiation can also be inhibited by its neighbor during subsequent transcription events. This supercoiling-mediated refractory period results in a loss of correlated transcription of both the divergent and tandem *GAL* genes. Moreover, we uncovered that in topoisomerase-deficient cells, *GAL10* is strongly inhibited by transcription-generated negative supercoils of its downstream *GAL7* gene, further reducing simultaneous transcription at the divergent *GAL1* and *GAL10* transcription. Overall, our data reveals that rapid supercoiling release is crucial to coordinate the transcription dynamics of neighboring eukaryotic genes.

In wildtype, transcription bursting of both the divergent and tandem gene pairs is coupled (**Figure 1**). Previous studies have shown similar co-expression of closely-positioned divergent gene pairs (*49*–*51*), which has fueled the prediction that negative supercoiling propagation induces correlated transcription at neighboring genes similar to bacteria (*21, 24, 52*). Although we cannot exclude the possibility that DNA supercoiling contributes to a small degree to the simultaneous initiation in wildtype, our data suggests that excess supercoiling at the *GAL* locus mostly impedes, rather than facilitates, initiation (**Figures 2, 3, S2**). For tandem genes, considering the supercoiling-mediated inhibition of *GAL7*, simultaneous initiation may be the most optimal scenario to balance the torsional stress generated at the locus. For divergent genes, however, we propose that coupled *GAL1-GAL10* initiation mostly originates from Gal4 binding to the shared upstream activating sequences. We have previously shown that fluctuations in Gal4 binding directly cause fluctuations in *GAL10* transcription (*53*). Based on this, we expect that once Gal4 binds, it simultaneously activates *GAL1* and *GAL10*. In addition, looping interactions or 3D proximity of the *GAL1-10* and *GAL7* promoters may facilitate correlated Gal4 binding also at the tandem *GAL* genes (*54, 55*). Correlated transcription factor binding has previously been observed for SRF at early immediate response genes and adjacent genes, resulting in their correlated transcription (*50*). Lastly, simultaneous transcription initiation of adjacent genes may be caused by long-distance activation of transcription factors, transcription factor clustering or transcription factor activity gradients (*49, 56*–*58*).

At the *GAL* locus, we find that both positive and negative supercoiling accumulation impede transcription, causing mutual inhibition at the tandem genes pair. The functional relevance of this mutual inhibition is underscored by an increase in yeast fitness when *GAL7* is relocated from the *GAL* locus to a different chromosome (*59*). In prokaryotes, the inhibitory role of positive supercoils is well-established (*10, 19*). Similar inhibition by positive supercoils occurs in eukaryotes (*60*) and is already weakly apparent in wildtype at highly transcribed genes (**Figure 1**). However, we find that the inhibitory effect of negative supercoils is more dominant (**Figure 5**). In line with this, in mouse embryonic stem cells, a transient accumulation of negative supercoils during base excision repair was recently found to inhibit transcription and upon release, cause increased noise fluctuations (*61*). The level of negative supercoiling thus requires careful regulation by topoisomerase, since low levels of negative supercoils enhance transcription (*4*–*6, 22, 43, 62*), but hypernegative supercoiling is inhibitory (*11, 63*–*65*).

How negative supercoils inhibit transcription is unclear. We do not observe large effects on nucleosomes, Gal4 binding or TBP binding, suggesting a step downstream of TBP is affected. Negative supercoils can cause the formation of alternative DNA structures such as Z-DNA, quadruplexes or DNA cruciform (*66*), which may inhibit the formation of the preinitiation complex. In addition, negative supercoils may affect transcription at a post-initiation step. We observed shorter-duration, lower-intensity and lower-frequency bursts, which would be inconsistent with prolonged polymerase stalling or slowed elongation rate, but could be the results of faster elongation or premature termination, possibly by the formation of R-loops (*11, 30, 67, 68*).

Unlike in bacteria, our data suggests that in budding yeast topoisomerases are not present at limiting concentrations for transcription dynamics: (i) the refractory period observed upon topoisomerase depletion is not observed (for *GAL1-GAL10*) or present only very weakly (for *GAL10-GAL7)* in wildtype cells (**Figure 1**); (ii) inhibition of *GAL7* transcription or addition of a spacer, which partly rescue the *GAL1-GAL10* correlation in topoisomerase-deficient conditions, have no effect in wildtype (**Figure 4**). This fundamental difference in supercoiling regulation between bacteria and yeast may be the result of differences in the topoisomerase enzymes, as bacterial topoisomerases are specialized for positive or negative supercoils, whereas eukaryotic topoisomerases relieve both (*69*). In addition, eukaryotes may buffer positive supercoils by nucleosomes (*28, 27*). Nevertheless, our data indicates eukaryotic topoisomerases are also not present in large excess since basal degradation of topoisomerase levels by only 25% already causes large transcriptional effects. Topoisomerase levels may be tightly controlled to ensure that DNA supercoiling accumulation remains at a level that is beneficial for transcription while limiting harmful effects. The weak valleys in the *GAL10-GAL7* CCF in wildtype (**Figure 1**) suggest that topoisomerase levels are at the tipping point of this balance.

In higher eukaryotes, transcription-generated negative supercoils contribute to cohesin extrusion and may therefore be crucial for the formation of topologically associating domains (TADs) (*32, 70, 71*). This mechanism assumes propagation of negative supercoils over much larger genomic distances than 1.5kb, the distance at which negative supercoils were initially thought to spread (*23, 72*). Even though mammalian genes are spaced much further apart than in yeast, supercoiling-dependent cohesin extrusion may cause supercoiling-effects from adjacent genes at a much larger distance. In higher eukaryotes, the accumulation of negative supercoils during nuclear processes, such as base excision repair (*61*), could influence the transcription of genes throughout the TAD. Overall, our single-cell live-cell approach highlights how efficient release of torsional stress is necessary to prevent inhibition of neighboring eukaryotic genes.

## Acknowledgements

We thank Fred van Leeuwen and Stefano G. Manzo for critical reading of the manuscript and members of the T.L.L lab for helpful feedback and suggestions. We thank Evelina Tutucci for sharing the 12xMS2V6 plasmid (Addgene plasmid 104390), Daniela Delneri for sharing the loxP mutant plasmid, Benjamin Albert and David Shore for sharing Top1-AID, Top2-AID W303 haploids, Steve Buratowski for sharing the TBP antibody and Joaquim Roca for sharing gyrase (pSTS77) and Topo I (YEptopA-PGAL1) plasmids. We thank the Research High Performance Computing Facility and the Genomics Core Facility of the NKI for assistance.

## Author contributions

Conceptualization: H.P.P., T.L.L.

Data curation: H.P.P.

Formal analysis: H.P.P., S.C., W.P., I.B., T.L.L.

Funding acquisition: I.B., T.L.L.

Investigation: H.P.P.

Methodology: H.P.P., T.L.L.

Software: H.P.P., S.C., W.P., I.B., T.L.L.

Supervision: T.L.L.

Visualization: H.P.P., T.L.L.

Writing – original draft: H.P.P., T.L.L.

Writing – review & editing: H.P.P., T.L.L., S.C., W.P., I.B.

## Data and materials availability

The sequencing (MNase-seq) data from this publication have been deposited to the GEO database https://www.ncbi.nlm.nih.gov/geo/ and assigned the identifier GSE196945. Microscopy data is available upon request. Software is available at https://github.com/Lenstralab/livecell and https://github.com/Lenstralab/smFISH.

## Materials and Methods

### Yeast strains and plasmids

Haploid yeast cells of BY4741 and BY4742 backgrounds were transformed and mated to obtain the BY4743 diploids listed in **Table S1**. 12xMS2V6 loops were integrated at 5’ *GAL1* with a PCR product containing loxP-kanMX-loxP and at 5’ *GAL7* with loxP2272-kanMX-loxP2272, a loxP mutant to prevent recombination with wildtype loxP sequence. The kanMX was excised with inducible CRE recombinase. Plasmids containing the MS2 and PP7 coat proteins, fused to mScarlet and GFPEnvy, respectively (pTL174 and pTL333), were digested with PacI and integrated at the *URA3* locus. Auxin-inducible degron tags at *TOP1* and *TOP2* were amplified from YTL738 or pTL398 and integrated at the endogenous loci. Plasmid containing OsTIR1 (pTL231) was digested with PacI and integrated at the *HIS3* locus. Gal4UASscr and spacer mutations were made using CRISPR/Cas9 (*74*). The spacer sequence included convergent ADH1t and CUT60t terminator sequences to prevent transcriptional interference. All integrations were checked with PCR and sequencing. Gyrase and Topo I were ectopically expressed from plasmids. Strains, plasmids and oligos used to construct the strains can be found in **Tables S1, S2 and S3**, respectively.

### Live-cell imaging and analysis

Live-cell imaging of transcription dynamics was performed as previously described in (Brouwer et al., 2020; Donovan et al., 2019) with minor modifications. Cells were grown at 30°C for at least 14 h in synthetic complete media supplemented with 2% raffinose. The cells were imaged after 30 min galactose induction at 30°C at mid-log (optical density, OD_600_ 0.2–0.4) on a coverslip with an agarose pad consisting of synthetic complete media and 2% galactose. For auxin treatment, cells were treated with galactose for 30 min and with 500 µM auxin for 15 min before imaging. For auxinole treatment, cells were grown for at least 14 h in 500 µM auxinole and induced with galactose for 30 min before imaging.

Imaging was performed on a setup consisting of an inverted microscope (Zeiss AxioObserver), an alpha Plan-Apochromat 100x 1.46NA oil objective, an sCMOS camera (Hamamatsu ORCA Flash 4v3) with a 475-570 nm dichroic (Chroma 59012bs), a 570 nm longpass beamsplitter (Chroma T565lpxr-UF1), an UNO Top stage incubator and objective heater (OKOlab) at 30°C, LED excitation at 470/24 nm and 550/15 nm (SpectraX, Lumencor) at 0.20% and 0.40% power with an ND2 filter, resulting in a 62 mW/cm^2^ and 413 mW/cm^2^ excitation intensity. Wide-field images of GFPEnvy and mScarlet signals were acquired sequentially to prevent spectral crosstalk. Images were recorded at 10s interval for 30 min, with 9 *z*-stacks (Δz 0.5 µm) and 200 ms exposure using the Micro-Manager software (*75*). For each condition, at least 3 replicate datasets were acquired with a total at least 100 cells.

For image analysis, the intensity calculation and tracking of the transcription sites was calculated as previously described in (*53*) using a custom Python script (https://github.com/Lenstralab/livecell). The images were maximum intensity projected and then corrected for *xy*-drift in the stage using an affine transformation. Cells were segmented using Otsu thresholding and watershedding. The intensity of the TS was calculated for each color by fitting a 2D Gaussian mask after local background subtraction as described previously (*76*). To detect the TSs, initial intensity thresholds of 9 and 7 standard deviations (SD) from the mean background was used for *PP7* and *MS2* signals, respectively. For frames where no TS was detected, a second fit was made in the vicinity of the initial detected spots using lower intensity thresholds of 6 and 4 SD from the mean background for *PP7* and *MS2*, respectively. If no TSs were detected in a frame after the second fit, the intensity was measured at the *xy*-coordinates of the previous frame. The tracking within each cell was inspected visually, and the endpoint of each trace was manually set at the last frame where a TS was visible. Dividing cells and cells in which TSs were not reliably detected were excluded from the analysis. Only the cells that exhibited both *PP7* and *MS2* signals were considered for analysis. Cells with only signal in one channel were inspected but exhibited insufficient coat protein levels in the other channel for reliable analysis. For each cell, the background was estimated by fitting a Lorentzian distribution to intensities measured at four points at a fixed distance from the TS in each frame in the same cell. The mean background was subtracted from the intensity trace to obtain background-subtracted intensity traces.

To determine the on and off periods, the fluorescence signal was binarized by setting a threshold that was a specific standard deviation above the *MS2* and *PP7* background intensities. To determine a binarization threshold that captured the correct bursting kinetics, the sum of squared residuals between the ACFs of the binary signals (range 1.0-5.0, steps of 0.25) and the ACF of the analog fluorescence signal between 10s and 100s was calculated. The minimal residual was found at threshold values of 2.75 and 4.50 standard deviations above background, with residuals of 0.0025 and 0.0010 for *MS2* and *PP7*, respectively. The burst intensity was measured for frames where the binarized signal was on. The burst duration and burst frequency (defined as inverse of time between bursts) were calculated from the binarized data, and bursts with a duration of a single frame were considered as errors from the binarization and were excluded. Reported error bars and significance was calculated from bootstrapping with 1000 repetitions.

### Computing correlation functions and transcriptional overlap

For each time trace, autocorrelation (ACF) and crosscorrelation functions (CCF) were computed as

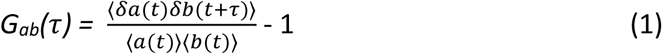

where <·> denotes the time average, *δa(t)* = *a(t) -* ⟨*a(t)*⟩and *a(t)* and *b(t)* can be combinations of the *MS2* and *PP7* time traces (*53, 76*). Correlation functions were computed using fast Fourier transforms and upon shifting the two signals, non-overlapping ends were trimmed. The functions were scaled for each trace individually. To correct for non-stationary effects (i.e. photobleaching, cell cycle, etc.), the global mean signal was used to calculate corrections, which were then subtracted. For single-trace correlation functions, each point was given a weight corresponding to the number of overlapping time intervals (τ) from the signals used in its computation. Correlation functions from single time traces were averaged together to reach statistical convergence. Bootstrapping was performed with 10,000 repetitions to obtain standard error of the mean correlation functions (SEM).

We used the ACFs and CCFs to calculate the normalized transcriptional overlap (called fractional overlap by Rodriguez *et al*.) (*36*), which provides an estimate of the fraction of bursts of one gene, which co-occur with the bursts of another gene. We normalized the cross-correlation functions of the *GAL1-GAL10* and *GAL10-GAL7* genes by their respective ACFs:

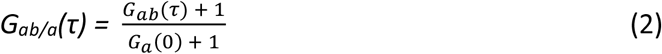

where *G*_*ab*_(*τ*) represents the CCFs of *GAL1-GAL10* or *GAL10-GAL7. G*_*a*_(0) represents the ACF amplitude at *τ*= 0 of *GAL10* in *GAL1-GAL10* or *GAL10-GAL7* pair. Each trace was normalized before the traces were averaged together.

To estimate the amplitude of the CCF at *τ* = 0, we fit the CCF with a Gaussian. The measured ACF amplitude at *τ* = 0 is overestimated due to shot noise, so to estimate the representative amplitudes, G_*a*_(0), we fit a line through the first 4 to 10 (omitting *τ* = 0) values of *G*_*a*_ and *G*_*b*_. The fit with the best coefficient of determination was used to extrapolate the values of *G*_*a*_(0).

Rodriguez *et al*. presented the transcriptional overlap calculation for a model with assumptions that the transcriptional events are square pulses of equal duration and height and are uniformly distributed over time. To confirm that this calculation can be applied for bursts with trapezoidal transcription events that are exponentially distributed, we simulated a 4-state model (ON-ON, ON-OFF, OFF-ON, OFF-OFF) for a gene pair where the promoter states are correlated, similar to *GAL1-GAL10*. We find that at lower transcription rates, the calculated normalized transcriptional overlap deviates from the theoretical values. However, for highly correlated gene pairs, as observed in the real data, the calculated transcriptional overlap from equation (2) matches the theoretical transcriptional overlap between the two genes.

### Single-molecule fluorescence *in situ* hybridization (smFISH) and analysis

Yeast cultures were grown to mid-log (OD_600_ 0.5) in 25 mL synthetic complete media with 2% raffinose and 2% galactose and smFISH was performed as previously described with minor modifications (*37, 53*). For auxinole treatment, cells were grown in synthetic complete media with 500 µM auxinole and 2% galactose. For the auxin timepoints, 100mL cultures were grown to OD_600_ 0.4 before being divided into 4×25 mL cultures and treated with 500 µM auxin for specified amount of time before fixation. If timepoint is not specified, cells were treated with auxin for 60 min. Cells were harvested at the same time after auxin addition to ensure the same OD.

Cells were fixed with 5% paraformaldehyde (Electron Microscopy Sciences, 15714-S) for 20 min, washed three times with buffer B (1.2 M sorbitol and 100 mM potassium phosphate buffer pH 7.5) and then spheroplasted with 300 units of lyticase (Sigma-Aldrich, L2524-25KU). Cells were then immobilized on poly-L-lysine-coated coverslips (Neuvitro) and permeabilized with 70% ethanol. Coverslips were hybridized for 4 h at 37°C with hybridization buffer containing 10% dextran sulfate, 10% formamide, 2×SSC, and 5 pmol of fluorescent probes. For FISH targeting the PP7 and MS2 repeats, four PP7 probes labeled with Quasar570 and 48 MS2 probes labeled with Quasar670 dyes were used. For FISH targeting *GAL1, GAL10* or *GAL7*, 48 probes labeled with Quasar570 (*GAL1* and *GAL7*) or Quasar670 (*GAL10*) were used (**Table S4**). Coverslips were washed 2× for 30 min with 10% formamide, 2×SSC at 37°C, then 1× with 2×SSC, and 1× for 5 min with PBS at room temperature. Coverslips were mounted on microscope slides using ProLong Gold mounting media with DAPI (Thermo Fisher, P36934).

Imaging was performed on two similar microscopes consisting of an inverted microscope (Zeiss AxioObserver), a Plan-Apochromat 40x 1.4NA oil DIC UV objective, a 1.60x optovar, and an sCMOS camera (Hamamatsu ORCA Flash 4v3). For Quasar570, a 562 nm longpass dichroic (Chroma T562lpxr), 595/50 nm emission filter (Chroma ET595/50m) and 550/15 nm LED excitation at full power (Spectra X, Lumencor) were used. For Quasar670, a 660 nm longpass dichroic (Semrock FF660-Di02-25×36 or Chroma T660lpxrxt), 697/60 nm emission filter (Chroma ET697/60m) and 640/30 nm LED excitation at full power (Spectra X, Lumencor) were used. For DAPI, either a 410nm/490nm/570nm/660nm dichroic (Chroma vcgr-spx-p01-PC), a 430/35 nm, 512/45 nm, 593/40 nm, 665 nm longpass emission filter (Chroma vcgr-spx-p01-EM) or a 425 nm longpass dichroic (Chroma T425lpxr) and a 460/50 nm emission filter (Chroma ET460/50m) and LED excitation at 395/25 nm at 25% power (Spectra X, Lumencor) were used. For each sample and each channel, we utilized the Micro-Manager software to acquire at least 50 fields-of-view, each consisting of a 21 *z*-stack (Δ*z* 0.3 µm) at 25 ms exposure for DAPI and 250 ms exposure for Quasar570 and Quasar670. For the smFISH experiments with the untagged topoisomerase-deficient haploids, all imaging settings were the same except a 1.25× optovar was used and each field-of-view consisted of 13 *z*-stack (Δ*z* 0.5 µm).

For smFISH image analysis, a custom-written Python script was used to detect, localize, and classify the spots (https://github.com/Lenstralab/smFISH). Cells and nuclei were segmented using Otsu thresholding and watershedding. Spots were localized by fitting a 3D Gaussian mask after local background subtraction (*76*). Cells in which no spots were detected were excluded from further analysis since a visual inspection indicated that these cells were not properly segmented or were improperly permeabilized. For each cell, the TS was defined as the brightest nuclear spot and the number of RNAs at each TS was determined by normalizing the intensity of each TS with the median fluorescent intensity of the cytoplasmic RNAs detected in all cells. Cells were further subclassified based on their cell cycle stage using the integrated DAPI intensity of each cell calculated from the maximum intensity projection images. A distribution of nuclear DAPI intensities was fit with a bimodal Gaussian model. The TS intensity was only analyzed in G1 cells, with nuclear intensities [1 SD, 0.75× SD] around the mean of the first peak.

Cells with fewer than 5 RNAs at the TS were classified as inactive, and cells with 5 or more RNAs at the TS were classified as active cells. Subsequently, the fraction of active cells for each gene and the Pearson correlation coefficients of the active cells were determined for various conditions. For smFISH experiments with *GAL10, GAL1*, and *GAL7* probes, spots were fit using 2D fitting, and the threshold to classify as an active cell was set to 2.5.

### Western blot and quantification

Yeast cultures were grown to mid-log (OD_600nm_ 0.4) in 25 mL synthetic complete media with 2% raffinose and 2% galactose. For auxinole treatment, cells were grown in synthetic complete media with 500 µM auxinole and 2% galactose. The cells were treated with 500 µM auxin for 15, 30, or 60 min. Cells were harvested at the same time to ensure the same OD. Cells were washed with PBS twice and then incubated in 200 mM NaOH for 10 min. The cells were pelleted and resuspended in 2× SDS-PAGE solvent (4% SDS, 20% glycerol, 0.1 M DTT, 0.125 M Tris-HCl pH 7.5 and Roche EDTA-free protease inhibitor cocktail) and boiled at 95°C for 5min. The lysates were centrifuged, the supernatant was collected and snap-frozen in liquid nitrogen and stored at −80°C.

To determine the loading volume, samples were first checked with a dot blot. The same wildtype control strain was used to ensure similar loading between experiments. For the western blot, samples were run on a 3-8% Tris-acetate gel (Thermo Fisher Scientific, EA0375PK2) at 100V for 2 hours and wet transferred (Bio-Rad, 1703930) on a nitrocellulose membrane at 300 mA for 4 hours. The membrane was washed with PBS for 5 min, blocked with 5% milk, dissolved in PBS, for 1 h at 18-22°C and incubated in 2% milk dissolved in TBS-T containing 1:1000 dilution of anti-cMyc (Thermo Fisher Scientific, #MA1-980) or anti-PGK (Thermo Fisher Scientific, #PA5-28612) primary antibodies, at 4°C for 14 hours. The membrane was washed with PBS for 5 min three times and incubated with 2% milk dissolved in TBS-T containing fluorescent anti-mouse (LI-COR, 926-32210) or anti-rabbit (LI-COR, 926-32211) secondary antibodies for 1 h at 18-22°C in the dark.

The fluorescence signal was quantified using ImageJ (*77*). A region of interest (ROI) was outlined around the largest sample and the same ROI was used for all samples on the membrane. The background was calculated by averaging the intensities of eight ROIs on the membrane where there was no signal present. The integrated intensities of the samples were background-subtracted and normalized to a no-OsTIR1 control strain to determine degradation upon OsTIR1 addition and auxin addition.

### Quantification of eGFP-tagged Topo I fluorescence

To quantify the Topo I-eGFP fluorescence, cells were segmented using Otsu thresholding and watershedding. The fluorescence of each pixel within the cell was integrated and subtracted by the mean background of the image. The background was defined as the pixels unoccupied by the cell masks. The background-subtracted intensities were normalized by the area of the cell.

### MNase-seq and analysis

Preparation and analysis of mono-nucleosomal DNA was performed as described previously (*53, 78*) with minor modifications and with two biological replicates. Haploid cells were grown in synthetic complete media with 2% raffinose or 2% galactose from OD_600_ 0.3 to OD_600_ 1.0, fixed in 1% paraformaldehyde, washed with 1 M sorbitol, treated with spheroplasting buffer (1M sorbitol, 1 mM β-mercaptoethanol, 10 mg/mL zymolyase 100T (US biological, Z1004.250)) and washed twice with 1 M sorbitol. Spheroplasted cells were treated with 0.01171875 or 0.1875 U micrococcal nuclease (Sigma-Aldrich, N5386-200UN) in digestion buffer (1 M sorbitol, 50 mM NaCl, 10 mM Tris pH7.4, 5 mM MgCl_2_, 0.075% NP-40, 1 mM β-mercaptoethanol, 0.5 mM spermidine) at 37°C. After 45 min, reactions were terminated on ice with 25 mM EDTA and 0.5% SDS. Samples were treated with proteinase K for 1 h at 37°C and decrosslinked overnight at 65°C. Digested DNA was extracted with phenol/chloroform (PCI 15:14:1), precipitated with NH_4_-Ac, and treated with 0.1 mg/mL RNaseA/T1. The extent of digestion was checked on a 3% agarose gel.

Sequencing libraries were prepared using the KAPA HTP Library Preparation Kit (07961901001, KAPA Biosystems) using 1 mg of input DNA, 5 mL of 10 mM adapter, double-sided size selection before and after amplification using 10 cycles. Adapters were created by ligation of the Universal adapter to individual sequencing adapters (**Table S5**). Libraries were checked on Bioanalyzer High Sensitivity DNA kit (Agilent) and sequencing was performed on a NextSeq550. Paired-end 2×75-bp reads were aligned to the reference genome SacCer3 using Bowtie 2. Nucleosome dyads were found by taking the middle of each paired read of insert size between 95 and 225 bp and were smoothed with a 31-bp window (*78*).

To determine the position of the +1 nucleosome for each gene, the coverage was determined in a 4000 bp window around the annotated TSS. For each gene, the coverage was summed and smoothed using a Gaussian filter with 40 bp window. The peaks were determined using a peak calling function. The +1 nucleosome was defined as the first peak after the minimum of the smoothed coverage. To compute the metagene plot, genes were aligned at the +1 nucleosome based on classifications in the unperturbed condition and the coverages were summed and normalized by the number of genes. Data are deposited in GEO with accession number: GSE196945.

### ChIP-qPCR

ChIP was performed as described in (*79*) using three biological replicates. Cells were diluted in the morning to OD_600_ 0.2 in 100 mL of synthetic complete media and grown for at least 2 doublings to an OD_600_ 0.8. Addition of auxin was staggered such that all time points were ready at the same OD (0.8). The cells were cross-linked for 5 min by addition of 37% formaldehyde (Sigma-Aldrich #252549) to a final concentration of 2%. The formaldehyde was quenched using a final concentration of 1.5 M of Tris (tris(hydroxymethyl)aminomethane) for 1 min. Subsequently, the cells were pelleted by centrifugation, and washed with TBS (150 mM NaCl, 10 mM Tris pH 7.5) twice and then snap-frozen in liquid nitrogen and stored at −80°C.

The cells were lysed by resuspending the pellets in FA lysis buffer (50 mM HEPES-KOH pH 7.5, 150 mM NaCl, 1 mM EDTA pH 8.0, 1% Triton X-100, 0.1% Na-deoxycholate, 0.1% SDS) containing the protease inhibitors aprotinin, pepstatin A, leupeptin and PMSF and then the cell walls were disrupted using 0.5 mm zirconium/silica beads (BioSpec Products, #11079105z) and bead beating 7×2 min in a Mini-Beadbeater-96 (Biospec #1001). The lysate was recovered and centrifuged to remove cell debris. The supernatant was subsequently fragmented by sonicating the samples for 10 cycles of 15 s on, 30 s off using a Bioruptor Pico sonicator (Diagenode #B01060010).

For the immunoprecipitation, the fragmented chromatin was incubated with 1 μg of anti-Gal4 antibody (Abcam #135397) or anti-TBP (from Steve Buratowski lab) antibody and then bound to magnetic beads (Dynabeads protein G, Life Technologies #10004D). The beads were washed twice with PBS and once with PBS-T and the cross-links were reversed by incubating in Tris-EDTA+1% SDS at 65°C for 14 h. The next morning, RNA was degraded with 0.005 mg/mL RNAse A/T1 (Thermo Scientific #EN0551), and proteins were digested with 0.04 mg/mL proteinase K (Roche #03115852001). DNA was recovered using a DNA purification kit (Bioline #52060). The binding levels of Gal4 and TBP were determined using qPCR. The protocol used the SensiFAST No-ROX kit (Bioline #98020) and a LightCycler 480 system for detection. The relative occupancy was calculated by estimating the input recovered in the IP.

### Rpb1-seq analysis

Annotated genes from YeastMine (https://yeastmine.yeastgenome.org/yeastmine/begin.do) were used to classify adjacent genes as divergent, tandem, or convergent. Gene pairs in the Rpb1-ChIP dataset that matched the YeastMine data were used to calculate the fold change. The vehicle condition was used to define the expression levels: low expression includes values 25^th^ percentile and below; intermediate expression includes 25^th^ to 75^th^ percentile; high expression includes 75^th^ percentile to 90^th^ percentile and very high expression includes the top 10 percentile.

For each gene pair orientation, pairs were binned based on the expression level of the upstream gene and the log_2_ fold change (log_2_(Rpb1 integrated reads in auxin treatment/Rpb1 integrated reads in vehicle)) was calculated of the downstream gene. The same approach was used to calculate the log_2_ fold change of the upstream gene with varied expression levels of the downstream gene. For divergent and convergent genes, the log_2_ fold changes of the upstream and downstream genes were combined into a single plot since the order is irrelevant.

