## Supplemental files for "DNA supercoiling restricts the transcriptional bursting of neighboring eukaryotic genes"

### Supplementary materials

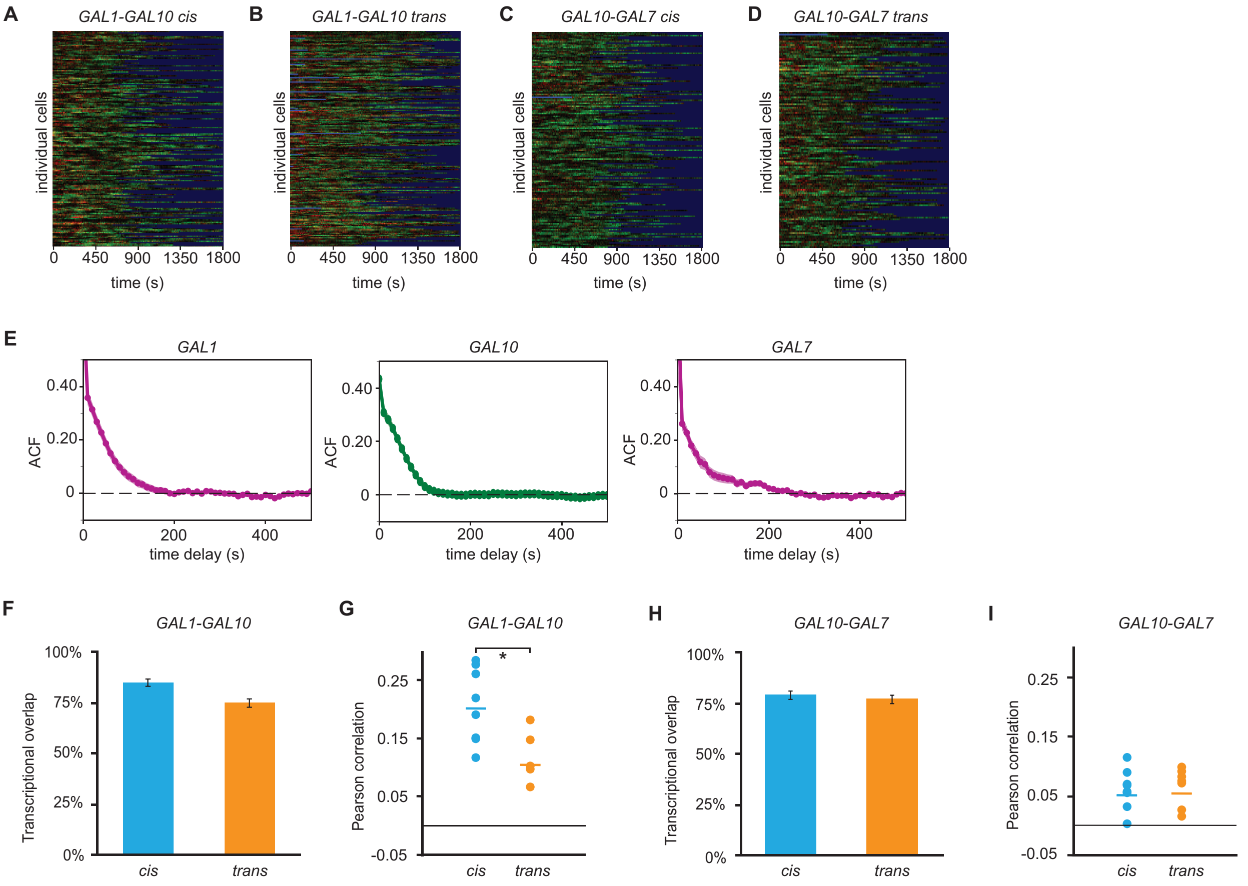


**Fig. S1: Transcriptional bursting of the divergent and tandem *GAL* genes is temporally coupled**

(**A-B**) Heatmaps of *GAL1* (red) and *GAL10* (green) TS intensities of individual cells (rows) for *cis* (*n* = 179 cells) and *trans* (*n* = 197 cells) configurations. Red and green indicate the intensities of *GAL1* and *GAL10* TSs, respectively. Blue indicates frames excluded from the analysis.

(**C-D**) Same as (**A-B**), for *GAL10* (green) and *GAL7* (red) for *cis* (*n* = 148 cells) and *trans* (*n* = 125 cells) configurations.

(**E**) Auto-correlation functions of *GAL1*, *GAL10* and *GAL7*. Shaded area indicates SEM.

(**F**) Transcriptional overlap of *cis*- and *trans*-labeled divergent *GAL1-GAL10* genes. Error bars indicate SEM.

(**G**) Pearson correlation coefficients of *GAL1-GAL10* nascent transcription by smFISH of *cis*- and *trans*-labeled genes. Each circle represents single replicate smFISH experiment (*n* = 7 for *cis* and *n* = 5 for *trans*). Horizontal lines represent mean. All experiments consist of at least 500 cells. *$p$<0.05, determined by two-tailed t-test.

(**H**) Same as (**F**) for tandem *GAL10-GAL7* genes.

(**I**) Same as (**G**) for tandem *GAL10-GAL7* genes (*n* = 6 for *cis* and *n* = 7 for *trans*).

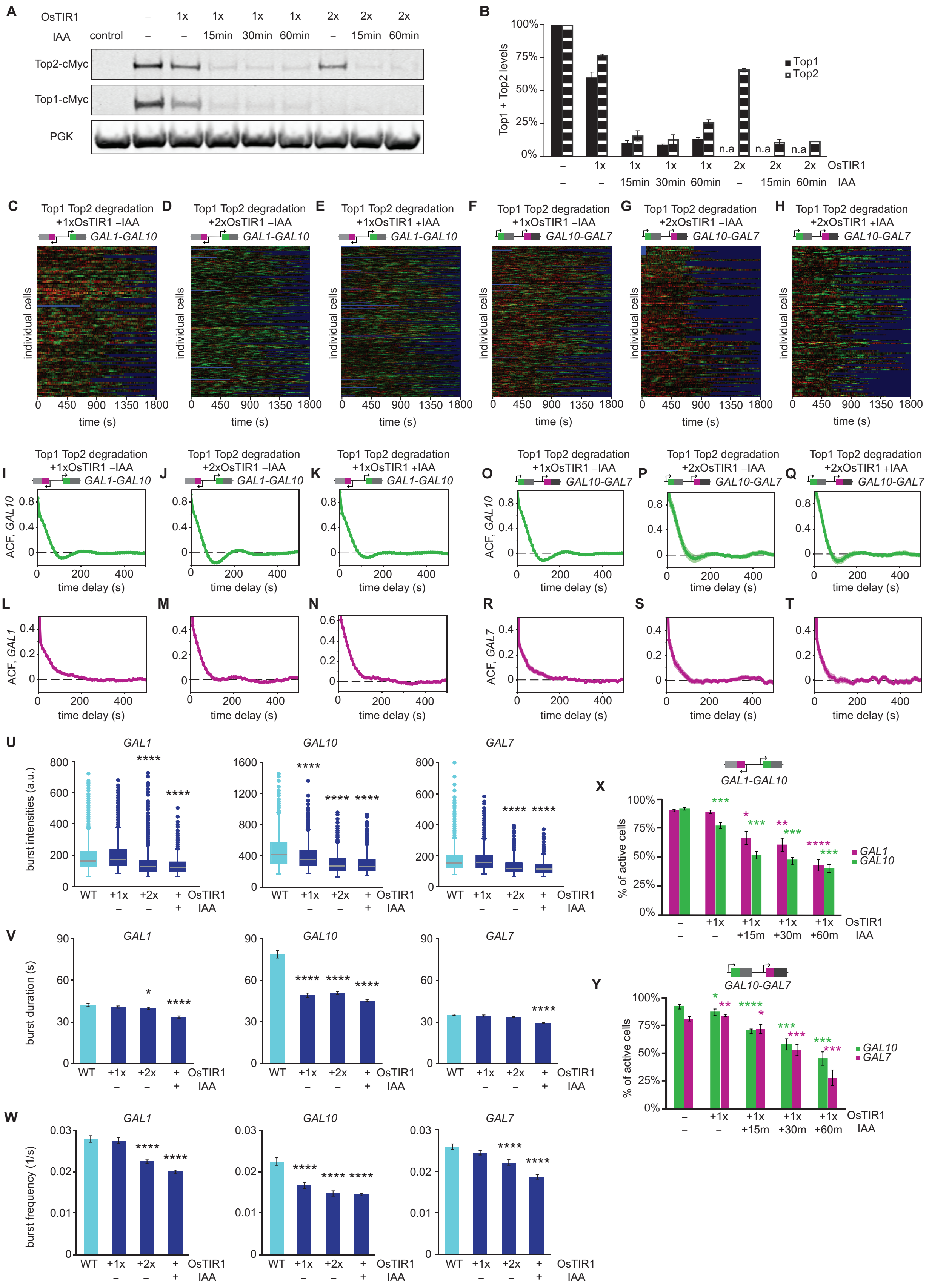

A
OsTIR1 – 1x 1x 1x 1x 2x 2x 2x
IAA control – – 15min 30min 60min – 15min 60min
Top2-cMyc
Top1-cMyc
PGK
B
Top1 + Top2 levels
100%
75%
50%
25%
0%
Top1
Top2
n.a
– 1x 1x 1x 1x 2x 2x 2x OsTIR1
– – 15min 30min 60min – 15min 60min IAA
C Top1 Top2 degradation +1xOsTIR1 –IAA GAL1-GAL10
D Top1 Top2 degradation +2xOsTIR1 –IAA GAL1-GAL10
E Top1 Top2 degradation +1xOsTIR1 +IAA GAL1-GAL10
F Top1 Top2 degradation +1xOsTIR1 –IAA GAL10-GAL7
G Top1 Top2 degradation +2xOsTIR1 –IAA GAL10-GAL7
H Top1 Top2 degradation +2xOsTIR1 +IAA GAL10-GAL7
individual cells
0 450 900 1350 1800
time (s)
I Top1 Top2 degradation +1xOsTIR1 –IAA GAL1-GAL10
J Top1 Top2 degradation +2xOsTIR1 –IAA GAL1-GAL10
K Top1 Top2 degradation +1xOsTIR1 +IAA GAL1-GAL10
O Top1 Top2 degradation +1xOsTIR1 –IAA GAL10-GAL7
P Top1 Top2 degradation +2xOsTIR1 –IAA GAL10-GAL7
Q Top1 Top2 degradation +2xOsTIR1 +IAA GAL10-GAL7
ACF, GAL10
time delay (s)
L
M
N
R
S
T
ACF, GAL1
ACF, GAL7
time delay (s)
U
GAL1
GAL10
GAL7
burst intensities (a.u.)
WT +1x +2x + OsTIR1
– – + IAA
V
burst duration (s)
W
burst frequency (1/s)
X
GAL1-GAL10
% of active cells
GAL1
GAL10
– +1x +1x +1x +1x OsTIR1
– – +15m +30m +60m IAA
Y
GAL10-GAL7
% of active cells
GAL10
GAL7
– +1x +1x +1x +1x OsTIR1
– – +15m +30m +60m IAA

**Fig. S2: Conditional degradation of topoisomerases results in refractory periods and reduced simultaneous initiation**

(**A**) Western blot of auxin-inducible degron-tagged Top1 and Top2, which are depleted rapidly and efficiently with one or two copies of OsTIR1, at various timepoints after auxin (IAA) addition. PGK is the loading control. Control strain is a wildtype diploid of BY4743 background.

(**B**) Quantification of the Top1 and Top2 levels in various conditions. n.a. indicates that the cells contained a degron sequence but did not express c-Myc for protein detection. $n$ = 3. Error bars indicate SEM.

(**C-E**) Heatmaps of the *GAL1* (red) and *GAL10* (green) TS intensities of individual cells (rows) with (C) basal degradation of topoisomerases from one copy of OsTIR1, (D) basal degradation of topoisomerases from two copies of OsTIR1, and (E) topoisomerase degradation 15 min post auxin (IAA) addition. $n$ = 117 cells, 214 cells, and 273 cells, respectively. Blue indicates frames excluded from the analysis.

(**F-H**) Same as (**C-E**) for *GAL10* (green) and *GAL7* (red). $n$ = 188 cells, 119 cells, and 143 cells.

(**I-N**) Auto-correlation functions of the divergent *GAL10* (I-K) and *GAL1* (L-N) in cells with (I,L) a basal degradation of topoisomerases from one copy of OsTIR1, (J,M) a basal degradation of topoisomerases from two copies of OsTIR1, and (K,N) topoisomerase degradation upon the addition of auxin (IAA). $n$ = 117 cells, 214 cells, and 273 cells, respectively. Shaded area indicates SEM.

(**O-T**) Same as (**I-N**) for the tandem *GAL10* (O-Q) and *GAL7* (R-T), $n$ = 188 cells, 119 cells, and 143 cells, respectively.

(**U**) Burst intensities, defined as background-subtracted TS intensities during on periods, for *GAL1* (left), *GAL10* (middle) and *GAL7* (right) in various topoisomerase-deficient conditions. The box indicates quartiles, the horizontal line inside the box indicates the median, and the whiskers indicate 1.5 times the interquartile range of the box. Data points outside the whiskers are indicated with circles. ****$p$<0.0001, determined by bootstrap hypothesis testing compared to wildtype.

(**V**) Burst durations for *GAL1* (left), *GAL10* (middle) and *GAL7* (right) in various topoisomerase-deficient conditions. *$p$<0.05; ****$p$<0.0001, determined by bootstrap hypothesis testing compared to wildtype.

(**W**) Burst frequencies, defined as the inverse of time between bursts, for *GAL1* (left), *GAL10* (middle) and *GAL7* (right) in various topoisomerase-deficient conditions. ****$p$<0.0001, determined by bootstrap hypothesis testing compared to wildtype.

(**X-Y**) Percentage of transcriptionally active cells, determined by smFISH, for (X) *GAL1* and *GAL10* genes and (Y) *GAL10* and *GAL7* genes. Errors bars indicate SEM. *$p$<0.05; **$p$<0.01; ***$p$<0.001; ****$p$<0.0001, determined by two-tailed t-test and compared to -OsTIR1 -IAA condition.

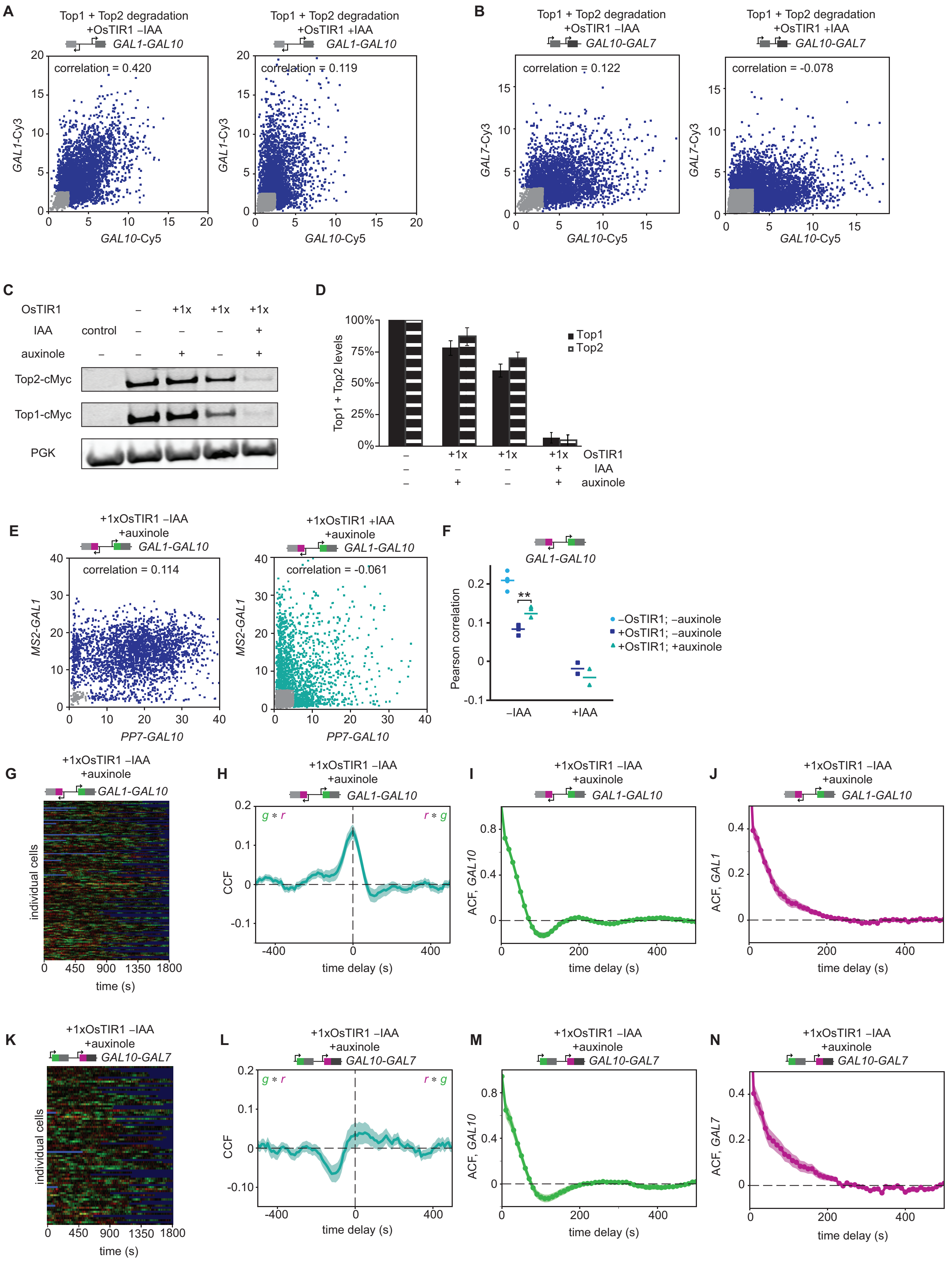
A
Top1 + Top2 degradation
+OsTIR1 –IAA
GAL1-GAL10
correlation = 0.420
GAL1-Cy3
GAL10-Cy5
Top1 + Top2 degradation
+OsTIR1 +IAA
GAL1-GAL10
correlation = 0.119
GAL1-Cy3
GAL10-Cy5
B
Top1 + Top2 degradation
+OsTIR1 –IAA
GAL10-GAL7
correlation = 0.122
GAL7-Cy3
GAL10-Cy5
Top1 + Top2 degradation
+OsTIR1 +IAA
GAL10-GAL7
correlation = -0.078
GAL7-Cy3
GAL10-Cy5
C
OsTIR1
IAA
auxinole
control
+1x
Top2-cMyc
Top1-cMyc
PGK
D
Top1 + Top2 levels
100%
75%
50%
25%
0%
Top1
Top2
OsTIR1
IAA
auxinole
E
+1xOsTIR1 –IAA
+auxinole
GAL1-GAL10
correlation = 0.114
MS2-GAL1
PP7-GAL10
+1xOsTIR1 +IAA
+auxinole
GAL1-GAL10
correlation = -0.061
MS2-GAL1
PP7-GAL10
F
GAL1-GAL10
Pearson correlation
**
–OsTIR1; –auxinole
+OsTIR1; –auxinole
+OsTIR1; +auxinole
–IAA
+IAA
G
+1xOsTIR1 –IAA
+auxinole
GAL1-GAL10
individual cells
time (s)
H
+1xOsTIR1 –IAA
+auxinole
GAL1-GAL10
g * r
r * g
CCF
time delay (s)
I
+1xOsTIR1 –IAA
+auxinole
GAL1-GAL10
ACF, GAL10
time delay (s)
J
+1xOsTIR1 –IAA
+auxinole
GAL1-GAL10
ACF, GAL1
time delay (s)
K
+1xOsTIR1 –IAA
+auxinole
GAL10-GAL7
individual cells
time (s)
L
+1xOsTIR1 –IAA
+auxinole
GAL10-GAL7
g * r
r * g
CCF
time delay (s)
M
+1xOsTIR1 –IAA
+auxinole
GAL10-GAL7
ACF, GAL10
time delay (s)
N
+1xOsTIR1 –IAA
+auxinole
GAL10-GAL7
ACF, GAL7
time delay (s)

**Fig. S3: Auxinole partially rescues the basal degradation of Top1 and Top2, as well as the associated phenotype**

(**A**) Scatterplots of the number of nascent transcripts at the *GAL1* and *GAL10* TSs determined by smFISH using *GAL1* and *GAL10* probes in haploids with degron-tagged Top1 and Top2. Cells without auxin (left, *n* = 4,104 cells) and after 60 min of auxin (right, *n* = 4,565 cells). Each datapoint represents a cell. Gray datapoints represent transcriptionally inactive cells that were excluded from the Pearson correlation coefficients analysis.

(**B**) Same as (**A**), for *GAL10* and *GAL7* in cells without auxin (left, *n* = 4,227 cells) and after 60 min of auxin (right, *n* = 11,638 cells).

(**C**) Western blot of Top1 and Top2 with auxinole treatment.

(**D**) Quantification of the Western blot for Top1 and Top2 levels with auxinole treatment. *n* = 2. Error bars indicate SEM.

(**E**) Scatterplots of the number of nascent transcripts at the *GAL1* and *GAL10* TSs, determined by smFISH, of cells treated with auxinole (left, n = 2,678 cells) or with both auxinole and auxin (middle, *n* = 3,199 cells) cells. Each datapoint represents a cell. Gray datapoints represent transcriptionally inactive cells that were excluded from the Pearson correlation coefficients calculation.

(**F**) Pearson correlation coefficient of *GAL1-GAL10* nascent transcription from smFISH in cells treated with auxinole and auxin. Each shape represents a single replicate smFISH experiment (*n* = 4, 3, 3, 2, 2 from left to right). Horizontal lines represent mean. All experiments consist of at least 500 cells. **$p$<0.01, determined by two-tailed t-test.

(**G,K**) Heatmaps of (G) *GAL1-GAL10* and (K) *GAL10-GAL7* TS intensities of individual cells (rows) with auxinole treatment. *n* = 143 cells and 77 cells, respectively. Red and green indicate the intensities of *GAL1* and *GAL10* TSs (G), or *GAL7* and *GAL10* (K), respectively. Blue indicates frames excluded from the analysis.

(**H,L**) *MS2-PP7* cross-correlation functions of (H) *GAL1-GAL10* and (L) and *GAL10-GAL7* after auxinole treatment. Shaded area indicates SEM.

(**I,M**) Auto-correlation function of *GAL10* from the (I) *GAL1-GAL10* gene pair and (M) and *GAL10-GAL7* gene pair after auxinole treatment. Shaded area indicates SEM.

(**J,N**) Similar to (**I, M**) for *GAL1* and *GAL7* from the (J) *GAL1-GAL10* gene pair and (N) and *GAL10-GAL7* gene pair, respectively, after auxinole treatment.

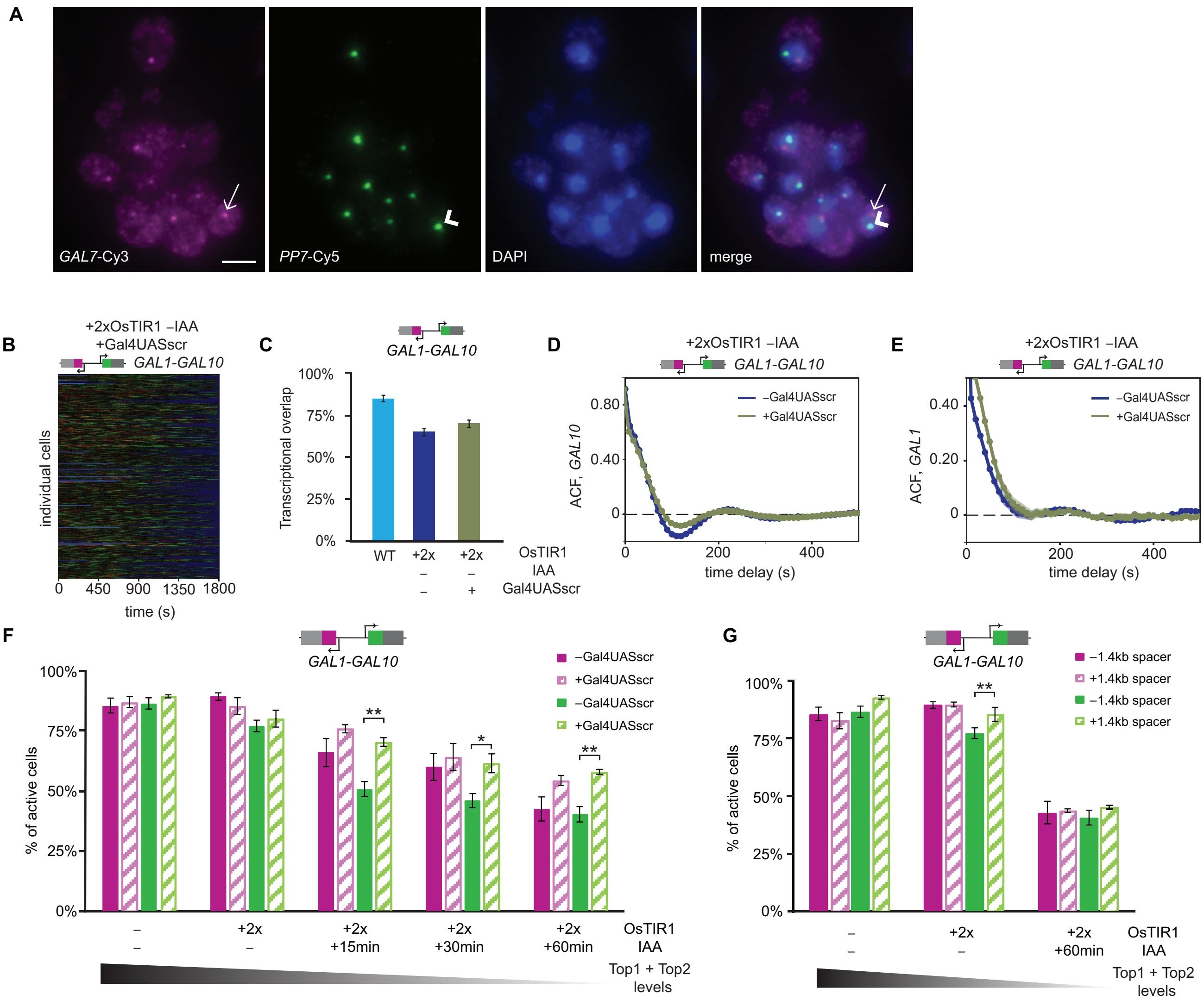


**Fig. S4: Transcription-generated supercoils from *GAL7* inhibit *GAL10* transcription in topoisomerase-deficient conditions**

(**A**) Example images of yeast cells with Gal4UASscr mutation at *GAL7* promoter to abolish expression. Cells were hybridized with fluorescent *GAL7*-Cy3 probes, *PP7*-Cy5 probes and DAPI. Each diploid cell has one *GAL7* TS (indicated by arrow), which does not colocalize with the *PP7* TS (indicated by arrowhead). Scale bar: 5 µm.

(**B**) Heatmap of *GAL1-GAL10* TS intensities of individual cells (rows) with mutation of Gal4UAS at the *GAL7* promoter. Red and green indicate the intensities of *GAL1* and *GAL10* TSs, respectively. Blue indicates frames excluded from the analysis. $n$ = 350 cells.

(**C**) Transcriptional overlap of the divergent *GAL1-GAL10* genes in wildtype (blue; same as **Figure S1C**), and topoisomerase-deficient cells without (navy, same as **Figure 3A**) and with (green) the Gal4UASscr mutation.

(**D-E**) Overlays of the ACFs for (D) *GAL10* and (E) *GAL1* to compare with and without Gal4UASscr mutation.

(**F**) Percentage of *GAL1* (magenta) and *GAL10* (green) active cells, determined by smFISH, in topoisomerase-deficient conditions without (solid bars) and with UASscr in the promoter of *GAL7* (patterned bars). Error bars indicate SEM. *$p<0.05$; **$p<0.01$, determined by two-tailed t-test.

(**G**) Same as (**F**) for cells without (solid bars) and with a spacer sequence between *GAL10* and *GAL7* (patterned bars). **$p<0.01$, determined by two-tailed t-test.

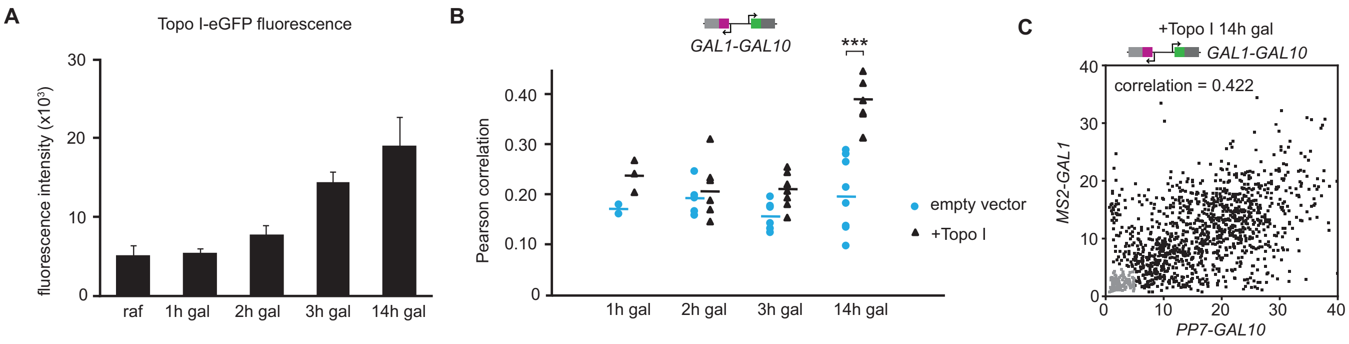


**Fig. S5: Transcription inhibition at the *GAL* locus is predominantly caused by negative supercoils**

(**A**) Quantification of the fluorescence of eGFP-tagged Topo I in raffinose or after galactose addition. Each condition consists of at least 20 cells.

(**B**) Pearson correlation coefficients *GAL1-GAL10* nascent transcription from smFISH at various timepoints post galactose addition. Each shape represents a single replicate smFISH experiment ($n$ = 2, 3, 5, 6, 5, 7, 8, 6 from left to right). All experiments consist of at least 500 cells. ***$p$<0.001, determined by two-tailed t-test.

(**C**) Scatterplot of the number of nascent transcripts at the *MS2-GAL1* and *PP7-GAL10* TSs, determined by smFISH, after 14 h of Topo I overexpression. $n$ = 1,320 cells.

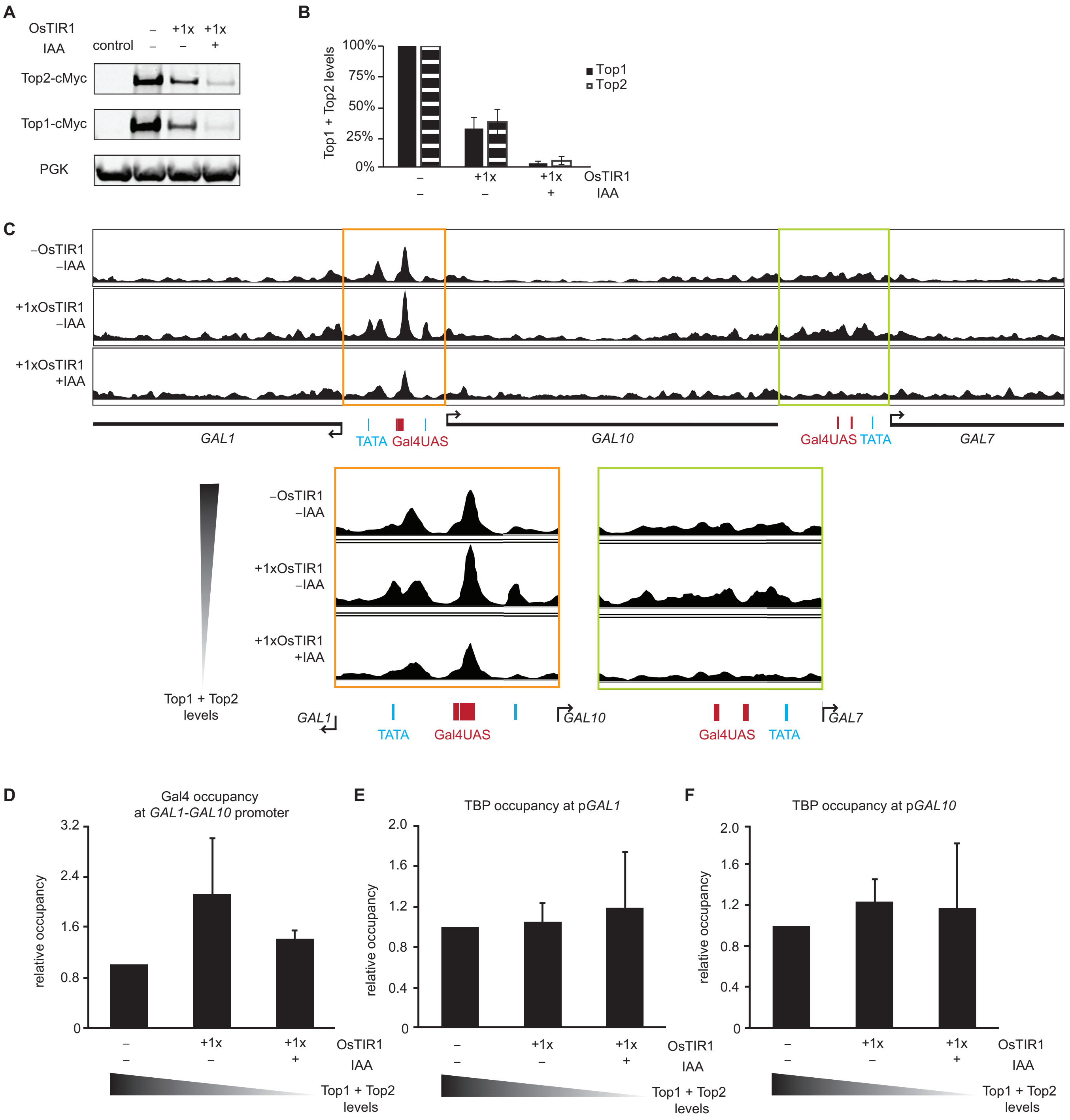


**Fig. S6: Supercoiling-mediated inhibition is not caused by altered fragile nucleosome positioning, or altered Gal4 or TBP occupancy**

(**A**) Western blot of degron-tagged Top1 and Top2 protein levels in haploid cells with or without 1x integration of OsTIR, in the presence or absence of auxin.

(**B**) Quantification of the Western blot for Top1 and Top2 levels. $n$ = 2. Error bars indicate SEM.

(**C**) MNase-seq profiles depicting the fragile nucleosome midpoint positions at the *GAL* locus in WT (top) and topoisomerase-deficient (middle, bottom) haploid cells. Zoom-in of the *GAL1-GAL10* promoter

region and *GAL10-GAL7* promoter regions depicted in orange and green boxes, respectively. Positions of Gal4 binding sites (magenta) and TATA boxes (blue) are indicated.
(**D-F**) Bar plots showing the relative (D) Gal4 occupancy at the *GAL1-GAL10* promoter
(**E**) TBP occupancy at the *GAL1* promoter and (F) TBP occupancy at the *GAL10* promoter in wildtype and topoisomerase-deficient cells, as obtained from ChIP experiments normalized over input, and normalized to wildtype binding. *n* = 3. Error bars indicate SEM.

**Table S1: Yeast strains used in this study**

| Yeast strain | Source |
|---|---|
| *S. cerevisiae BY4743: MATa/α his3Δ1/his3Δ1 leu2Δ0/leu2Δ0 LYS2/lys2Δ0 met15Δ0/MET15 ura3Δ0/ura3Δ0* | Euroscarf |
| *YTL047: BY4742 with 14xPP7 5' GAL10* | Donovan et al. 2019 |
| *yTL738: W303 background, MATa HIS3+, ADE+, trp1-1, leu2-3 pADH1-OsTIR1:URA3 and TOP1-AID-kanMX and TOP2-9myc-AID*-HPH* | Albert et al. 2019 |
| *yTL870: BY4743 with 14xPP7 5'GAL10/GAL10 and 12xMS2V6 5'GAL1/GAL1* | this study |
| *yTL885: BY4743 with 14xPP7 5'GAL10/GAL10 and GAL1/12xMS2V6 5'GAL1* | this study |
| *yTL988: BY4743 with 14xPP7 5'GAL10/GAL10 and 12xMS2V6 5'GAL1/GAL1 and ura3-pRPL15A-PCP-GFPEnvy/ura3-pPAB1-MCP-mScarlet* | this study |
| *yTL985: BY4743 with 14xPP7 5'GAL10/GAL10 and GAL1/12xMS2V6 5'GAL1 and ura3-pRPL15A-PCP-GFPEnvy/ura3-pPAB1-MCP-mScarlet* | this study |
| *yTL1000: BY4743 with 14xPP7 5'GAL10/GAL10 and 12xMS2V6 5'GAL1/GAL1 and his3-pADH1-OsTIR1/his3-pADH1-OsTIR1 and TOP1-AID-kanMX/TOP1-AID-kanMX and TOP2-AID*-9myc-HPH/TOP2-AID*-9myc-HPH* | this study |
| *yTL1090: BY4743 with 14xPP7 5'GAL10/GAL10 and 12xMS2V6 5'GAL7/GAL7* | this study |
| *yTL1091: BY4743 with 14xPP7 5'GAL10/GAL10 and 12xMS2V6 5'GAL7/GAL7* | this study |
| *yTL1096: BY4743 with 14xPP7 5'GAL10/GAL10 and 12xMS2V6 5'GAL7/GAL7 and ura3-pRPL15A-PCP-GFPEnvy/ura3-pPAB1-MCP-mScarlet* | this study |
| *yTL1097: BY4743 with 14xPP7 5'GAL10/GAL10 and GAL7/12xMS2V6 5'GAL7 and ura3-pRPL15A-PCP-GFPEnvy/ura3-pPAB1-MCP-mScarlet* | this study |
| *yTL1210: BY4743 with 14xPP7 5'GAL10/GAL10 and 12xMS2V6 5'GAL1/GAL1 and Gal4UASscr 5'GAL7/GAL7 and his3-pADH1-OsTIR1/his3-pADH1-OsTIR1 and TOP1-AID-kanMX/TOP1-AID-kanMX and TOP2-AID*-9myc-HPH/TOP2-AID*-9myc-HPH* | this study |
| *yTL1232: BY4743 with 14xPP7 5'GAL10/GAL10 and 12xMS2V6 5'GAL7/GAL7 and his3-pADH1-OsTIR1/his3-pADH1-OsTIR1 and TOP1-AID-kanMX/TOP1-AID-kanMX and TOP2-AID*-9myc-HPH/TOP2-AID*-9myc-HPH* | this study |
| *yTL1245: BY4742 with TOP1-AID-9myc-kanMX/TOP1-AID-9myc-kanMX and TOP2-AID*-9myc-HPH/TOP2-AID*-9myc-HPH* | this study |
| *yTL1248: BY4743 with 14xPP7 5'GAL10/GAL10 and 12xMS2V6 5'GAL1/GAL1 and pTL258 (pGAL1_GyrA-GyrB_URA)* | this study |
| *yTL1250: BY4743 with 14xPP7 5'GAL10/GAL10 and 12xMS2V6 5'GAL1/GAL1 and his3-pADH1-OsTIR1/his3-pADH1-OsTIR1 and TOP1-AID-kanMX/TOP1-AID-kanMX and TOP2-AID*-9myc-HPH/TOP2-AID*-9myc-HPH and pTL258 (pGAL1_GyrA-GyrB_URA)* | this study |
| *yTL1282: BY4743 with 14xPP7 5'GAL10/GAL10 and 12xMS2V6 5'GAL1/GAL1 and Gal4UASscr 5'GAL7/GAL7* | this study |
| *yTL1290: BY4742 with his3-pADH1-OsTIR1 and TOP1-AID-9myc-kanMX/TOP1-AID-9myc-kanMX and TOP2-AID*-9myc-HPH/TOP2-AID*-9myc-HPH* | this study |

| | |
|---|---|
| *yTL1292: BY4743 with 14xPP7 5'GAL10/GAL10 and 12xMS2V6 5'GAL1/GAL1 and TOP1-AID-9myc-kanMX/TOP1-AID-9myc-kanMX and TOP2-AID*-9myc-HPH/TOP2-AID*-9myc-HPH* | this study |
| *yTL1294: BY4743 with 14xPP7 5'GAL10/GAL10 and 12xMS2V6 5'GAL1/GAL1 and his3-pADH1-OsTIR1/his3Δ1 and TOP1-AID-kanMX/TOP1-AID-kanMX and TOP2-AID*-9myc-HPH/TOP2-AID*-9myc-HPH* | this study |
| *yTL1299: BY4743 with 14xPP7 5'GAL10/GAL10 and 12xMS2V6 5'GAL1/GAL1 and his3-pADH1-OsTIR1/his3Δ1 and TOP1-AID-9myc-kanMX/TOP1-AID-9myc-kanMX and TOP2-AID*-9myc-HPH/TOP2-AID*-9myc-HPH and ura3-pRPL15A-PCP-GFPEnvy/ura3-pPAB1-MCP-mScarlet* | this study |
| *yTL1300: BY4743 with 14xPP7 5'GAL10/GAL10 and 12xMS2V6 5'GAL7/GAL7 and his3-pADH1-OsTIR1/his3Δ1 and TOP1-AID-9myc-kanMX/TOP1-AID-9myc-kanMX and TOP2-AID*-9myc-HPH/TOP2-AID*-9myc-HPH and ura3-pRPL15A-PCP-GFPEnvy/ura3-pPAB1-MCP-mScarlet* | this study |
| *yTL1313: BY4743 with 14xPP7 5'GAL10/GAL10 and 12xMS2V6 5'GAL1/GAL1 and Gal4UASscr 5'GAL7/GAL7 and his3-pADH1-OsTIR1/his3-pADH1-OsTIR1 and TOP1-AID-kanMX/TOP1-AID-kanMX and TOP2-AID*-9myc-HPH/TOP2-AID*-9myc-HPH and ura3-pRPL15A-PCP-GFPEnvy/ura3-pPAB1-MCP-mScarlet* | this study |
| *yTL1347: BY4743 with 14xPP7 5'GAL10/GAL10 and 12xMS2V6 5'GAL1/GAL1 and pTL439 (pGAL1_topA_URA)* | this study |
| *yTL1348: BY4743 with 14xPP7 5'GAL10/GAL10 and 12xMS2V6 5'GAL1/GAL1 and his3-pADH1-OsTIR1/his3-pADH1-OsTIR1 and TOP1-AID-kanMX/TOP1-AID-kanMX and TOP2-AID*-9myc-HPH/TOP2-AID*-9myc-HPH and pTL439 (pGAL1_topA_URA)* | this study |
| *yTL1406: BY4743 with 14xPP7 5'GAL10/GAL10 and 12xMS2V6 5'GAL1/GAL1 and 388bp spacer 5'GAL7/GAL7 and TOP1-AID-9myc-kanMX/TOP1-AID-9myc-kanMX and TOP2-AID*-9myc-HPH/TOP2-AID*-9myc-HPH* | this study |
| *yTL1408: BY4743 with 14xPP7 5'GAL10/GAL10 and 12xMS2V6 5'GAL1/GAL1 and 1388bp spacer 5'GAL7/GAL7 and TOP1-AID-9myc-kanMX/TOP1-AID-9myc-kanMX and TOP2-AID*-9myc-HPH/TOP2-AID*-9myc-HPH* | this study |
| *yTL1455: BY4743 with 14xPP7 5'GAL10/GAL10 and 12xMS2V6 5'GAL1/GAL1 and 388bp spacer 5'GAL7/GAL7 and his3-pADH1-OsTIR1/his3-pADH1-OsTIR1 and TOP1-AID-9myc-kanMX/TOP1-AID-9myc-kanMX and TOP2-AID*-9myc-HPH/TOP2-AID*-9myc-HPH* | this study |
| *yTL1456: BY4743 with 14xPP7 5'GAL10/GAL10 and 12xMS2V6 5'GAL1/GAL1 and 1388bp spacer 5'GAL7/GAL7 and his3-pADH1-OsTIR1/his3-pADH1-OsTIR1 and TOP1-AID-9myc-kanMX/TOP1-AID-9myc-kanMX and TOP2-AID*-9myc-HPH/TOP2-AID*-9myc-HPH* | this study |
| *yTL1625: BY4743 with 14xPP7 5'GAL10/GAL10 and 12xMS2V6 5'GAL1/GAL1 and pTL523 (pGAL1_topA-linker-eGFP_URA)* | this study |
| *yTL1633: BY4743 with 14xPP7 5'GAL10/GAL10 and 12xMS2V6 5'GAL1/GAL1 and 500bp spacer 5'GAL7/GAL7 and his3-pADH1-OsTIR1/his3-pADH1-OsTIR1* | this study |

| | |
|---|---|
| *and TOP1-AID-9myc-kanMX/TOP1-AID-9myc-kanMX and TOP2-AID*-9myc-HPH/TOP2-AID*-9myc-HPH* | |
| *yTL1634: BY4743 with 14xPP7 5'GAL10/GAL10 and 12xMS2V6 5'GAL1/GAL1 and 700bp spacer 5'GAL7/GAL7 and his3-pADH1-OsTIR1/his3-pADH1-OsTIR1 and TOP1-AID-9myc-kanMX/TOP1-AID-9myc-kanMX and TOP2-AID*-9myc-HPH/TOP2-AID*-9myc-HPH* | this study |
| *yTL1635: BY4743 with 14xPP7 5'GAL10/GAL10 and 12xMS2V6 5'GAL1/GAL1 and 900bp spacer 5'GAL7/GAL7 and his3-pADH1-OsTIR1/his3-pADH1-OsTIR1 and TOP1-AID-9myc-kanMX/TOP1-AID-9myc-kanMX and TOP2-AID*-9myc-HPH/TOP2-AID*-9myc-HPH* | this study |
| *yTL1636: BY4743 with 14xPP7 5'GAL10/GAL10 and 12xMS2V6 5'GAL1/GAL1 and 1100bp spacer 5'GAL7/GAL7 and his3-pADH1-OsTIR1/his3-pADH1-OsTIR1 and TOP1-AID-9myc-kanMX/TOP1-AID-9myc-kanMX and TOP2-AID*-9myc-HPH/TOP2-AID*-9myc-HPH* | this study |

**Table S2: Plasmids used in this study**

| Plasmid | Source |
|---|---|
| pTL014: pGAL1_CRErecombinase_URA | Lenstra et al. 2015 |
| pTL031: 14xPP7 with loxP-kanMX-loxP | Donovan et al. 2019 |
| pTL131: pML104 Cas9 guide RNA construct | Laughery et al. 2015 |
| pTL174: SIVURA_pRPL15A_PCP-NLS-GFPEnvy | this study |
| pTL181: pUC 12xMS2V6_U-var_50nt_linker loxP-kanMX-loxP | gift from Evelina Tutucci |
| pTL253: 1kb spacer centered with convergent ADH1t (5'; scrambled antisense TATA) and CUT60t (3') | this study |
| pTL258: pGAL1_GyrA-GyrB_URA | gift from Joaquim Roca |
| pTL327: pGPD_CRE-EBD78_URA | gift from Fred van Leeuwen |
| pTL333: SIVURA_pPAB1_MCP-NLS_mScarlet | this study |
| pTL337: pFA6a-lox2272-natNT2-lox2272 | gift from Daniela Delneri |
| pTL338: pUC 12xMS2V6_U-var_50nt_linker lox2272-kanMX-lox2272 | this study |
| pTL361: pML104 with gRNA to GAL10-GAL7 intergenic region | this study |
| pTL355: pML104 with gRNA to Gal4UAS distal to GAL7 | this study |
| pTL365: pML104 with gRNA to Gal4UAS proximal to GAL7 | this study |
| pTL398: TOP1-AID-9myc-kanMX | this study |
| pTL439: pGAL1_topA_URA | gift from Joaquim Roca |
| pTL523: pGAL1_topA-linker-eGFP_URA | this study |

**Table S3: Oligos used in this study**

| Oligo | Source |
|---|---|
| 12xMS2V6-loxP-kanMX-loxP tag at GAL1_F:<br>gtatcaacaaaaaattgttaatatacctctatactttaacgtcaaggagaaaaaactataccgctctagaactagtggatcc | IDT |
| 12xMS2V6-loxP-kanMX-loxP tag at GAL1_R:<br>ttcctttgcgctagaattgaactcaggtacaatcacttcttctgaatgagatttagtcatgcataggccactagtggatctg | IDT |

| | |
|---|---|
| 12xMS2V6-loxP2272-kanMX-loxP2272 tag at GAL7_F: gggcattattatgcagagcatcaacatgataaaaaaaaacagttgaatattccctcaaaaccgctctagaactagtggatcc | IDT |
| 12xMS2V6-loxP2272-kanMX-loxP2272 tag at GAL7_R: ggttagtggattgtaacgtctatgggaatggctagaaaaatcaaattcttcagcagtcatgcataggccactagtggatctg | IDT |
| TOP1-AID-kanMX tag_F: gaagccaccattcacaagag | IDT |
| TOP1-AID-kanMX tag_R: attggaggaagcatggatgc | IDT |
| TOP1-AID-9myc-kanMX tag_F: tcgacgaaaagaaacaagaattcgaaaag | IDT |
| TOP1-AID-9myc-kanMX tag_R: aatgcgaacttgatgcgtgaatgtatttgcttctcccctatgctgcgtttctttgcgttacagtatagcgaccagcattcac | IDT |
| TOP2-AID*-9myc-HPH tag_F: aacaggacgatgtagccact | IDT |
| TOP2-AID*-9myc-HPH tag_R: accaggcatggagcttatct | IDT |
| Gal4UAS proximal to GAL7 repair: tccttaacccaaaaataagggaaagggtccaaaaagcgctccgaattgctcgacgagtgatccgaaggactggctatacagtgttcacaaa atagcc | IDT |
| Gal4UAS distal to GAL7 repair: ttcttaaattgctttgcctctccttttggaaagctatacttgcagtccatggagggtcaaggctcattagatatattttctgtcattttcctt aacccaa | IDT |
| 388bp spacer at GAL7 promoter repair_F: caaaaattgaaaatctatggaaagatatggacggtagcaacaagaatatagcacgagccgtgaggcgcgccacttctaaa | IDT |
| 388bp spacer at GAL7 promoter repair_R: tttcactgaagcgcttcgcaatagttgtgagtgatatcaaaagtaacgaaatgaactctgagaagactataagctaaaaatgtagacaac | IDT |
| 500bp spacer at GAL7 promoter repair_F: caaaaattgaaaatctatggaaagatatggacggtagcaacaagaatatagcacgagccgttcagctgcaataatatcggtatttattatgg | IDT |
| 500bp spacer at GAL7 promoter repair_R: tttcactgaagcgcttcgcaatagttgtgagtgatatcaaaagtaacgaaatgaactctggtttagcaatttcattttcagtaattcctg | IDT |
| 700bp spacer at GAL7 promoter repair_F: caaaaattgaaaatctatggaaagatatggacggtagcaacaagaatatagcacgagccgaaatcttcgatataaaaaatgcggtaatttcag | IDT |
| 700bp spacer at GAL7 promoter repair_R: tttcactgaagcgcttcgcaatagttgtgagtgatatcaaaagtaacgaaatgaactctgggaacaccgtttggaattcttatatgg | IDT |
| 900bp spacer at GAL7 promoter repair_F: caaaaattgaaaatctatggaaagatatggacggtagcaacaagaatatagcacgagccgtggaggttgacgactaggca | IDT |
| 900bp spacer at GAL7 promoter repair_R: tttcactgaagcgcttcgcaatagttgtgagtgatatcaaaagtaacgaaatgaactctggtctctgtgtgtgtacggatg | IDT |
| 1100bp spacer at GAL7 promoter repair_F: caaaaattgaaaatctatggaaagatatggacggtagcaacaagaatatagcacgagccggttgtcacagctaatattccattttgaattgac | IDT |
| 1100bp spacer at GAL7 promoter repair_R: tttcactgaagcgcttcgcaatagttgtgagtgatatcaaaagtaacgaaatgaactctgtcgacgaattaccatgaaaagctca | IDT |
| 1388bp spacer at GAL7 promoter repair_F: tttcactgaagcgcttcgcaatagttgtgagtgatatcaaaagtaacgaaatgaactctgccacttagaccggaatgtca | IDT |
| 1388bp spacer at GAL7 promoter repair_R: caaaaattgaaaatctatggaaagatatggacggtagcaacaagaatatagcacgagccgggtcactggagtattagcca | IDT |
| primer 1.1 Illumina: aatgatacggcgaccaccgagat | IDT |
| primer 2.1 Illumina: caagcagaagacggcatacga | IDT |

**Table S4: smFISH probes used in this study**

| smFISH probe | Source |
|---|---|
| PP7-1: [Cy3]atatcgtctgctcctttcta | IDT |
| PP7-2: [Cy3]atatgctctgctggtttcta | IDT |
| PP7-3: [Cy3]gcaattaggtaccttaggat | IDT |
| PP7-4: [Cy3]aatgaacccgggaatactgc | IDT |
| 12xMS2V6-1: [Quasar-670]cgcaagcgagagtgaagacg | Biosearch Technologies |
| 12xMS2V6-2: [Quasar-670]acacagggtattcctctgtg | Biosearch Technologies |
| 12xMS2V6-3: [Quasar-670]gaatgttctttgtagcaccg | Biosearch Technologies |
| 12xMS2V6-4: [Quasar-670]catgtgacactggtgagcaa | Biosearch Technologies |
| 12xMS2V6-5: [Quasar-670]aaggtttcctgactcgcttg | Biosearch Technologies |
| 12xMS2V6-6: [Quasar-670]tcgggataatggtgcgatgc | Biosearch Technologies |
| 12xMS2V6-7: [Quasar-670]ggtaatcctgcgtgtcgatt | Biosearch Technologies |
| 12xMS2V6-8: [Quasar-670]tttgacggggaacagagtgt | Biosearch Technologies |
| 12xMS2V6-9: [Quasar-670]tgcatgacgtaagccgactc | Biosearch Technologies |
| 12xMS2V6-10: [Quasar-670]tgcaccatgcatgcggtaat | Biosearch Technologies |
| 12xMS2V6-11: [Quasar-670]acgagtagacatgccgatat | Biosearch Technologies |
| 12xMS2V6-12: [Quasar-670]ccacacgagtagatttgact | Biosearch Technologies |
| 12xMS2V6-13: [Quasar-670]atagaagtggtcggtagtcc | Biosearch Technologies |
| 12xMS2V6-14: [Quasar-670]cctgcacaaccgaaaagatg | Biosearch Technologies |
| 12xMS2V6-15: [Quasar-670]cttctcgtacatcggcagaa | Biosearch Technologies |
| 12xMS2V6-16: [Quasar-670]ggtagtcgagaagcgtaatc | Biosearch Technologies |
| 12xMS2V6-17: [Quasar-670]atctgcacaccatgtatgat | Biosearch Technologies |
| 12xMS2V6-18: [Quasar-670]gtattcctccattggcaaaa | Biosearch Technologies |
| 12xMS2V6-19: [Quasar-670]catgggttggttgacagaga | Biosearch Technologies |
| 12xMS2V6-20: [Quasar-670]gccatagcagagtgtaaact | Biosearch Technologies |
| 12xMS2V6-21: [Quasar-670]ttttggctcgcagggtattc | Biosearch Technologies |
| 12xMS2V6-22: [Quasar-670]cgcaaggcagatgcaataca | Biosearch Technologies |
| 12xMS2V6-23: [Quasar-670]cgtattcgtccctgtgaatg | Biosearch Technologies |
| 12xMS2V6-24: [Quasar-670]agagcacccatattgaagta | Biosearch Technologies |
| 12xMS2V6-25: [Quasar-670]aaatggtcctgatgacgacg | Biosearch Technologies |
| 12xMS2V6-26: [Quasar-670]atatatgcgcggtagtcctg | Biosearch Technologies |
| 12xMS2V6-27: [Quasar-670]aagtatccgcacgagtgctg | Biosearch Technologies |
| 12xMS2V6-28: [Quasar-670]cgcgtaacaataggaatccc | Biosearch Technologies |
| 12xMS2V6-29: [Quasar-670]cagagctcggtattcctgag | Biosearch Technologies |
| GAL1-yeast-1: [Quasar-570]gtgacataagaaccgtccaa | Biosearch Technologies |
| GAL1-yeast-2: [Quasar-570]ctaaagccctgtttcttatt | Biosearch Technologies |
| GAL1-yeast-3: [Quasar-570]cataatgttctgcaacgacc | Biosearch Technologies |
| GAL1-yeast-4: [Quasar-570]tgtagaactcattggcaagg | Biosearch Technologies |
| GAL1-yeast-5: [Quasar-570]tttaaacggagtagccttca | Biosearch Technologies |
| GAL1-yeast-6: [Quasar-570]gtgatcttagggtacttgac | Biosearch Technologies |

| | |
|---|---|
| GAL1-yeast-7: [Quasar-570]gtaaaaccgagaagtcacaa | Biosearch Technologies |
| GAL1-yeast-8: [Quasar-570]cactgccagttggtacatca | Biosearch Technologies |
| GAL1-yeast-9: [Quasar-570]tctttgttaaccgttcgatg | Biosearch Technologies |
| GAL1-yeast-10: [Quasar-570]atttcttaattatgctcggg | Biosearch Technologies |
| GAL1-yeast-11: [Quasar-570]aatccggtttagcatcataa | Biosearch Technologies |
| GAL1-yeast-12: [Quasar-570]ttgaactcaggtacaatcac | Biosearch Technologies |
| GAL1-yeast-13: [Quasar-570]ggtaattcctttgcgctaga | Biosearch Technologies |
| GAL1-yeast-14: [Quasar-570]agattcagaatacacatgct | Biosearch Technologies |
| GAL1-yeast-15: [Quasar-570]aatgctggtttagagacgat | Biosearch Technologies |
| GAL1-yeast-16: [Quasar-570]gggcggtttcaaacttgtta | Biosearch Technologies |
| GAL1-yeast-17: [Quasar-570]tctaccaggcgatctagcaa | Biosearch Technologies |
| GAL1-yeast-18: [Quasar-570]ggacatatgataaccagggc | Biosearch Technologies |
| GAL1-yeast-19: [Quasar-570]tttgcagctgtagtgactt | Biosearch Technologies |
| GAL1-yeast-20: [Quasar-570]aaccaagtgaacagtacaac | Biosearch Technologies |
| GAL1-yeast-21: [Quasar-570]aaatcgaacttcctttgagc | Biosearch Technologies |
| GAL1-yeast-22: [Quasar-570]tttaccttttctatgttgcc | Biosearch Technologies |
| GAL1-yeast-23: [Quasar-570]agtcattaatttcacagcct | Biosearch Technologies |
| GAL1-yeast-24: [Quasar-570]attcaatatcgccgttccag | Biosearch Technologies |
| GAL1-yeast-25: [Quasar-570]gaaatgttgtgatatctggc | Biosearch Technologies |
| GAL1-yeast-26: [Quasar-570]aagagtgagcaacatggaga | Biosearch Technologies |
| GAL1-yeast-27: [Quasar-570]tccggtgcaagtttctttag | Biosearch Technologies |
| GAL1-yeast-28: [Quasar-570]gttcatcaaggcaccaaatt | Biosearch Technologies |
| GAL1-yeast-29: [Quasar-570]cattttctagctcagcatca | Biosearch Technologies |
| GAL1-yeast-30: [Quasar-570]attcatatagacagctgccc | Biosearch Technologies |
| GAL1-yeast-31: [Quasar-570]caatctctggacaagaacat | Biosearch Technologies |
| GAL1-yeast-32: [Quasar-570]agattacctttattcgtgct | Biosearch Technologies |
| GAL1-yeast-33: [Quasar-570]atttgacaaagcaatggaac | Biosearch Technologies |
| GAL1-yeast-34: [Quasar-570]aacaaagctaatttcatggt | Biosearch Technologies |
| GAL1-yeast-35: [Quasar-570]agtagtctcttgtgaattct | Biosearch Technologies |
| GAL1-yeast-36: [Quasar-570]cagaaagtaaaacaacaccg | Biosearch Technologies |
| GAL1-yeast-37: [Quasar-570]taccgccattgttaacacca | Biosearch Technologies |
| GAL1-yeast-38: [Quasar-570]gcgagaacaattcaaggatt | Biosearch Technologies |
| GAL1-yeast-39: [Quasar-570]tcaaacgggaaccatatgat | Biosearch Technologies |
| GAL1-yeast-40: [Quasar-570]ttcctcaccgcaaacagag | Biosearch Technologies |
| GAL1-yeast-41: [Quasar-570]cagaggagcactggcaaa | Biosearch Technologies |
| GAL1-yeast-42: [Quasar-570]cttttcggccaatggtct | Biosearch Technologies |
| GAL1-yeast-43: [Quasar-570]ttcgtcggcagtaaagctc | Biosearch Technologies |
| GAL1-yeast-44: [Quasar-570]ctttaagacttgaaatctcact | Biosearch Technologies |
| GAL1-yeast-45: [Quasar-570]acaagggtgttcgcaat | Biosearch Technologies |
| GAL1-yeast-46: [Quasar-570]cagaagacttgcagcccg | Biosearch Technologies |
| GAL1-yeast-47: [Quasar-570]cgagagactcttcaactagta | Biosearch Technologies |
| GAL1-yeast-48: [Quasar-570]gcaacggcacaaatgaat | Biosearch Technologies |

| | |
|---|---|
| GAL7-yeast-1: [Quasar-570]gcagacaagaaatcaccggt | Biosearch Technologies |
| GAL7-yeast-2: [Quasar-570]aaaaccttgtcttgcttcgt | Biosearch Technologies |
| GAL7-yeast-3: [Quasar-570]atcaatgtatctaccaggct | Biosearch Technologies |
| GAL7-yeast-4: [Quasar-570]ctttccagttctcttgattg | Biosearch Technologies |
| GAL7-yeast-5: [Quasar-570]aatcttggacccgtaagttt | Biosearch Technologies |
| GAL7-yeast-6: [Quasar-570]tcgctgtgacacttttatca | Biosearch Technologies |
| GAL7-yeast-7: [Quasar-570]aacaattcggtccctacatg | Biosearch Technologies |
| GAL7-yeast-8: [Quasar-570]atttttgatgtctccatggt | Biosearch Technologies |
| GAL7-yeast-9: [Quasar-570]tcgtgctatattcttgttgc | Biosearch Technologies |
| GAL7-yeast-10: [Quasar-570]actgaagcgcttcgcaatag | Biosearch Technologies |
| GAL7-yeast-11: [Quasar-570]gaaaaattaccccctctact | Biosearch Technologies |
| GAL7-yeast-12: [Quasar-570]tccgaagtatagctttccaa | Biosearch Technologies |
| GAL7-yeast-13: [Quasar-570]cccttatttttgggttaagg | Biosearch Technologies |
| GAL7-yeast-14: [Quasar-570]ttcagcttggctattttgtg | Biosearch Technologies |
| GAL7-yeast-15: [Quasar-570]acctgcttttatatctttgc | Biosearch Technologies |
| GAL7-yeast-16: [Quasar-570]tcatgttgatgctctgcata | Biosearch Technologies |
| GAL7-yeast-17: [Quasar-570]acgtctatgggaatggctag | Biosearch Technologies |
| GAL7-yeast-18: [Quasar-570]ccatgaatcggttagtggat | Biosearch Technologies |
| GAL7-yeast-19: [Quasar-570]tgacctaaccaaggtctttt | Biosearch Technologies |
| GAL7-yeast-20: [Quasar-570]tcatacaatggagctgtggg | Biosearch Technologies |
| GAL7-yeast-21: [Quasar-570]ctcttttgttaccaggacat | Biosearch Technologies |
| GAL7-yeast-22: [Quasar-570]tatcttgggtttaggttacc | Biosearch Technologies |
| GAL7-yeast-23: [Quasar-570]cctaacggcagcataatcat | Biosearch Technologies |
| GAL7-yeast-24: [Quasar-570]cctcattggaatcattctgt | Biosearch Technologies |
| GAL7-yeast-25: [Quasar-570]acaattgcctctcacagatt | Biosearch Technologies |
| GAL7-yeast-26: [Quasar-570]ctggagagatcgtcagtcaa | Biosearch Technologies |
| GAL7-yeast-27: [Quasar-570]gcttatgattttctcttgct | Biosearch Technologies |
| GAL7-yeast-28: [Quasar-570]ccatgtggatgtaagttgga | Biosearch Technologies |
| GAL7-yeast-29: [Quasar-570]cttcactagggatggattct | Biosearch Technologies |
| GAL7-yeast-30: [Quasar-570]actcttgacttctctcttga | Biosearch Technologies |
| GAL7-yeast-31: [Quasar-570]aaggtctcaaatggccagat | Biosearch Technologies |
| GAL7-yeast-32: [Quasar-570]cccattgagtatgggaaact | Biosearch Technologies |
| GAL7-yeast-33: [Quasar-570]actcaattcatcaccagtcg | Biosearch Technologies |
| GAL7-yeast-34: [Quasar-570]gctgatctcagtaaaggtgg | Biosearch Technologies |
| GAL7-yeast-35: [Quasar-570]ctttgaggctcacctaacaa | Biosearch Technologies |
| GAL7-yeast-36: [Quasar-570]tagtttttcagcagcttgtt | Biosearch Technologies |
| GAL7-yeast-37: [Quasar-570]ccactttctttttacagtct | Biosearch Technologies |
| GAL7-yeast-38: [Quasar-570]attcaggcattagtgctttt | Biosearch Technologies |
| GAL7-yeast-39: [Quasar-570]gtgtcagatgtcacagtttc | Biosearch Technologies |
| GAL7-yeast-40: [Quasar-570]tgctgcaacatccaattagg | Biosearch Technologies |
| GAL7-yeast-41: [Quasar-570]aacataggtgcaggatttcc | Biosearch Technologies |
| GAL7-yeast-42: [Quasar-570]cactgctcttctgcataatt | Biosearch Technologies |

| | |
|---|---|
| GAL7-yeast-43: [Quasar-570]tcgatgccatctttattcac | Biosearch Technologies |
| GAL7-yeast-44: [Quasar-570]gagttgctagttgcaacaca | Biosearch Technologies |
| GAL7-yeast-45: [Quasar-570]tagcttccattgcgttatta | Biosearch Technologies |
| GAL7-yeast-46: [Quasar-570]tctgaagatcctgtagggag | Biosearch Technologies |
| GAL10-yeast-1: [Quasar670]aggagtcttcaacctgcaa | Biosearch Technologies |
| GAL10-yeast-2: [Quasar670]aaaggattctcagtagtcca | Biosearch Technologies |
| GAL10-yeast-3: [Quasar670]tggcctcgacaccccttaa | Biosearch Technologies |
| GAL10-yeast-4: [Quasar670]catatcttcagcggaaaatc | Biosearch Technologies |
| GAL10-yeast-5: [Quasar670]atagtcacaaatcttgcgtc | Biosearch Technologies |
| GAL10-yeast-6: [Quasar670]ggcttgaaatctggtgccg | Biosearch Technologies |
| GAL10-yeast-7: [Quasar670]tcactttcaggtcaacaatg | Biosearch Technologies |
| GAL10-yeast-8: [Quasar670]atagccaagaacaactgatt | Biosearch Technologies |
| GAL10-yeast-9: [Quasar670]gaattcgacaggttatcagc | Biosearch Technologies |
| GAL10-yeast-10: [Quasar670]caaatacccttcctcatttt | Biosearch Technologies |
| GAL10-yeast-11: [Quasar670]gcgcctatataagcactatc | Biosearch Technologies |
| GAL10-yeast-12: [Quasar670]catacctgccgatcgtg | Biosearch Technologies |
| GAL10-yeast-13: [Quasar670]actaaacttacccttcgaaa | Biosearch Technologies |
| GAL10-yeast-14: [Quasar670]acggttaactgatagtcttt | Biosearch Technologies |
| GAL10-yeast-15: [Quasar670]ctatgattcgcattaacgcc | Biosearch Technologies |
| GAL10-yeast-16: [Quasar670]tctgtggaaagaaccgatac | Biosearch Technologies |
| GAL10-yeast-17: [Quasar670]aaggattttgaatgatgggt | Biosearch Technologies |
| GAL10-yeast-18: [Quasar670]cctggctacagaatcataag | Biosearch Technologies |
| GAL10-yeast-19: [Quasar670]atgtactcggcggtaaaa | Biosearch Technologies |
| GAL10-yeast-20: [Quasar670]tcggtgtccttctcattatc | Biosearch Technologies |
| GAL10-yeast-21: [Quasar670]ttaccaatagatcacctgga | Biosearch Technologies |
| GAL10-yeast-22: [Quasar670]ttgggcaacgttcacagtat | Biosearch Technologies |
| GAL10-yeast-23: [Quasar670]ccagcagtcaatttaccttt | Biosearch Technologies |
| GAL10-yeast-24: [Quasar670]ttaaatttattggcgtcgct | Biosearch Technologies |
| GAL10-yeast-25: [Quasar670]cctcaatagtgtctccatat | Biosearch Technologies |
| GAL10-yeast-26: [Quasar670]ttgaacgcaccataatctcc | Biosearch Technologies |
| GAL10-yeast-27: [Quasar670]attacccgtaggaatcatgt | Biosearch Technologies |
| GAL10-yeast-28: [Quasar670]ggctttgtagagttaaaggt | Biosearch Technologies |
| GAL10-yeast-29: [Quasar670]acaatcaaactggggatttt | Biosearch Technologies |
| GAL10-yeast-30: [Quasar670]ttgacttggcttagcatttt | Biosearch Technologies |
| GAL10-yeast-31: [Quasar670]aacctcatagaagggaatgt | Biosearch Technologies |
| GAL10-yeast-32: [Quasar670]atcgggatgaaaagccttga | Biosearch Technologies |
| GAL10-yeast-33: [Quasar670]tggctctgtacttaaaact | Biosearch Technologies |
| GAL10-yeast-34: [Quasar670]cagcagacaagaaatcaccg | Biosearch Technologies |
| GAL10-yeast-35: [Quasar670]accttgtcttgcttcgtaac | Biosearch Technologies |
| GAL10-yeast-36: [Quasar670]aatgtatctaccaggctcaa | Biosearch Technologies |
| GAL10-yeast-37: [Quasar670]ctttgtaactgagctgtcat | Biosearch Technologies |
| GAL10-yeast-38: [Quasar670]acacaatctttccagttctc | Biosearch Technologies |

| | |
|---|---|
| GAL10-yeast-39: [Quasar670]ccttttcggtcacacaaatc | Biosearch Technologies |
| GAL10-yeast-40: [Quasar670]caatcttggacccgtaagtt | Biosearch Technologies |
| GAL10-yeast-41: [Quasar670]agtgaattaccgaatcaatttta | Biosearch Technologies |
| GAL10-yeast-42: [Quasar670]cacctacagcctttaaacca | Biosearch Technologies |
| GAL10-yeast-43: [Quasar670]gtgatagtatctcagcggga | Biosearch Technologies |
| GAL10-yeast-44: [Quasar670]cctgtaaccaaaacaattttaga | Biosearch Technologies |
| GAL10-yeast-45: [Quasar670]ctaataaaacgacagttccc | Biosearch Technologies |
| GAL10-yeast-46: [Quasar670]atttggaaacgttgtattgt | Biosearch Technologies |
| GAL10-yeast-47: [Quasar670]caccatagacagtagcagaa | Biosearch Technologies |
| GAL10-yeast-48: [Quasar670]atttgggaatctcgtagcat | Biosearch Technologies |

**Table S5: Sequencing adapters used in this study**

| **Sequencing adapter** | **Source** |
|---|---|
| Universal adapter:<br>aatgatacggcgaccaccgagatctacactctttccctacacgacgctcttccgatc*t | IDT |
| Sequencing adapter 1:<br>[5'Phos]gatcggaagagcacacgtctgaactccagtcac**gatcag**atctcgtatgccgtcttctgcttg | IDT |
| Sequencing adapter 2:<br>[5'Phos]gatcggaagagcacacgtctgaactccagtcac**tagctt**atctcgtatgccgtcttctgcttg | IDT |
| Sequencing adapter 3:<br>[5'Phos]gatcggaagagcacacgtctgaactccagtcac**ggctac**atctcgtatgccgtcttctgcttg | IDT |
| Sequencing adapter 4:<br>[5'Phos]gatcggaagagcacacgtctgaactccagtcac**cttcga**atctcgtatgccgtcttctgcttg | IDT |
| Sequencing adapter 5:<br>[5'Phos]gatcggaagagcacacgtctgaactccagtcac**agtcaa**atctcgtatgccgtcttctgcttg | IDT |
| Sequencing adapter 6:<br>[5'Phos]gatcggaagagcacacgtctgaactccagtcac**agttcc**atctcgtatgccgtcttctgcttg | IDT |
| Sequencing adapter 7:<br>[5'Phos]gatcggaagagcacacgtctgaactccagtcac**atgtca**atctcgtatgccgtcttctgcttg | IDT |
| Sequencing adapter 8:<br>[5'Phos]gatcggaagagcacacgtctgaactccagtcac**ccgtcc**atctcgtatgccgtcttctgcttg | IDT |
| Sequencing adapter 9:<br>[5'Phos]gatcggaagagcacacgtctgaactccagtcac**gtccgc**atctcgtatgccgtcttctgcttg | IDT |
| Sequencing adapter 10:<br>[5'Phos]gatcggaagagcacacgtctgaactccagtcac**gtgaaa**atctcgtatgccgtcttctgcttg | IDT |
| Sequencing adapter 11:<br>[5'Phos]gatcggaagagcacacgtctgaactccagtcac**cgtacg**atctcgtatgccgtcttctgcttg | IDT |
| Sequencing adapter 12:<br>[5'-Phos]gatcggaagagcacacgtctgaactccagtcac**attcct**atctcgtatgccgtcttctgcttg | IDT |

**Video S1: Example movie of *MS2-GAL1* (magenta) and *PP7-GAL10* (green).** A representative video of an induced yeast cell showing *MS2* (magenta) and *PP7* (green) transcription sites (TSs), which are outlined by the white boxes. *MS2* and *PP7* TSs were imaged sequentially at 10s intervals for 180 frames with 9 z-slices ($\Delta z$ = 0.5 $\mu$m). Video is a maximum intensity *xy*-projection with *xz*- and *yz*-projections shown as side views.
